# Targeting cellular DNA damage responses in cancer: An *in vitro*-calibrated agent-based model simulating monolayer and spheroid treatment responses to ATR-inhibiting drugs

**DOI:** 10.1101/841270

**Authors:** Sara Hamis, James Yates, Mark AJ Chaplain, Gibin G Powathil

## Abstract

We combine a systems pharmacology approach with an agent-based modelling approach to simulate LoVo cells subjected to AZD6738, an ATR (ataxia telangiectasia mutated and rad3-related kinase) inhibiting anti-cancer drug that can hinder tumour proliferation by targeting cellular DNA damage responses. The agent-based model used in this study is governed by a set of empirically observable rules. By adjusting only the rules when moving between monolayer and multi-cellular tumour spheroid simulations, whilst keeping the fundamental mathematical model and parameters intact, the agent-based model is first parameterised by monolayer *in vitro* data and is thereafter used to simulate treatment responses in *in vitro* tumour spheroids subjected to dynamic drug delivery. Spheroid simulations are subsequently compared to *in vivo* data from xenografts in mice. The spheroid simulations are able to capture the dynamics of *in vivo* tumour growth and regression for approximately eight days post tumour injection.

Translating quantitative information between *in vitro* and *in vivo* research remains a scientifically and financially challenging step in preclinical drug development processes. However, well-developed in *silico* tools can be used to facilitate this in *vitro* to in vivo translation, and in this article we exemplify how data-driven, agent-based models can be used to bridge the gap between *in vitro* and *in vivo* research. We further highlight how agent-based models, that are currently underutilised in pharmaceutical contexts, can be used in preclinical drug development.

## 1 Introduction

### 1.1 Bridging *in vitro* and *in vivo* research

Mathematical models, and their corresponding *in silico* tools, can be used to simulate both *in vitro* and in *vivo* scenarios that involve cancer cell populations, or tumours, and their responses to anti-cancer treatments (Rockne et al., 2019; Bruno et al., 2020; Stephanou et al., 2018; Brady-Nicholls et al., 2020; Scott et al., 2019). However, cancer cells in an *in vitro* cell culture experience a microenvironment that is significantly different from the microenvironment experienced by cancer cells in a solid tumour in vivo. As these microenvironments influence cell proliferation and the delivery of oxygen, drug and nutrient molecules to cells, it follows that the dynamics of a cancer cell population in vitro differs from the dynamics of a solid tumour in vivo. Consequently, translating data obtained by in vitro experiments into quantitative information that can guide or predict in vivo experiments remains a challenging, but important, step in drug development processes. As an intermediate step between monolayer cell cultures and in vivo tumours, multicellular tumour spheroids (in this study referred to as spheroids) provide in *vitro* models that are able to capture certain key-features of *in vivo* tumours such as intratumoural heterogeneity resulting from nutrient-gradients and resource-limited tumour growth (Nunes et al., 2019).

Agent-based models (ABMs) are used in many applications in mathematical biology but are underutilised in the context of pharmaceutical drug development (Cosgrove et al., 2015). An ABM consists of multiple, distinct agents that may interact with each other and their microenvironment. There exists different types of ABMs. For example, agents can be deformable or of fix size, and agent movements and neighbourhood-interactions can be constrained by an underlying lattice geometry (on-lattice models) or not (off-lattice models). Combining ABMs with hybrid modelling techniques allows for the integration of discrete and continuous variables describing tumour dynamics on multiple scales. A thorough review on various types of hybrid ABMs used to simulate tumour growth is provided by Rejniak and Anderson (Rejniak and Anderson, 2011). Furthermore, a number of open-source in *silico* tools, such as Chaste (Mirams et al., 2013), CompuCell3D (Swat et al., 2012) and PhysiCell (Ghaffarizadeh et al., 2018), are freely available to facilitate the implementation of ABMs.

In this study, we introduce a novel modelling approach that uses an agent-based mathematical model to bridge the gap between *in vitro* monolayer and spheroid research as a step towards bridging the gap between *in vitro* and *in vivo* research, as is conceptually illustrated in Figure 1. For a broader scope discussion on how to develop, calibrate and validate mathematical models that can predict novel anti-cancer therapies, we refer the reader to a recent article by Brady and Enderling (Brady and Enderling, 2019). In the ABM at the core of this modelling approach, an agent consists of one cancer cell or a group of cancer cells, where the behaviour and fate of each agent is governed by a set of empirically observable and well-established *modelling rules* that incorporate both intracellular and microenvironmental dynamic variables, as is described throughout Section 2. To account for differences between monolayer and spheroid scenarios, the modelling rules are adjusted when moving between monolayer and spheroid simulations. By only adjusting the rules, whilst keeping the fundamental mathematical model and parameters intact, when moving between monolayer and spheroid simulations, the mathematical framework can first be parameterised by monolayer data and thereafter be used to simulate spheroid treatment responses. To exemplify this modelling approach, we here simulate LoVo (human colon carcinoma) cells subjected to the anti-cancer drug AZD6738. The ABM is first calibrated by monolayer *in vitro* data and is thereafter used to simulate *in vitro* spheroids subjected to dynamic drug delivery. Spheroid simulations are subsequently compared to xenograft *in vivo* data. The *in vitro* and *in vivo* data used in this study are gathered from previous work by Checkley *et al*. (Checkley et al., 2015). The ABM used in this study is an extension of a model introduced by Powathil *et al*. (Powathil et al., 2012b).

**Figure 1:**
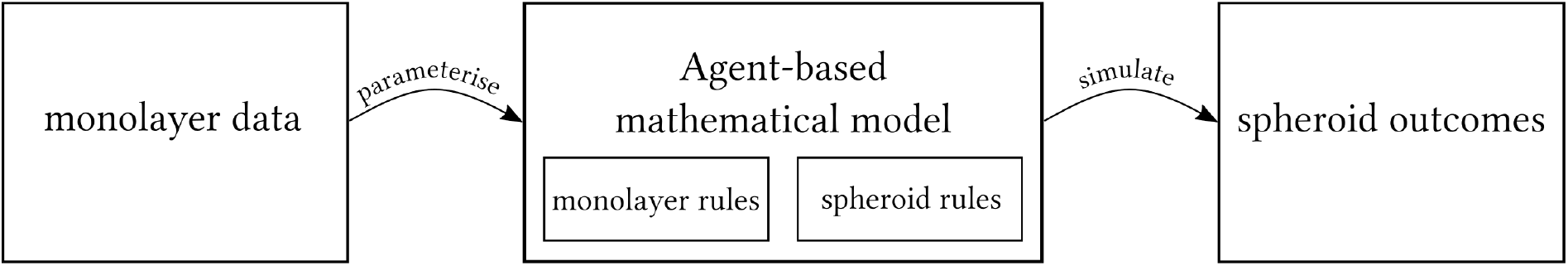
A schematic of the mathematical modelling approach used in this study. An agent-based mathematical model, that distinguishes between *in vitro* monolayer and spheroid modelling rules, is formulated. The mathematical model is first parameterised by *in vitro* monolayer data and is thereafter used to simulate spheroid dynamics.

### 1.2 DNA damage response inhibiting drugs

The deoxyribonucleic acid (DNA) in human cells is perpetually exposed to, potentially harmful, influences that can be derived from both exogenous and endogenous sources and events (Minchom et al., 2018; Sundar et al., 2017). Exogenous sources include ultraviolet radiation, ionising radiation and chemotherapeutic drugs, whilst erroneous DNA replication is an example an endogenous event yielding DNA damage (Minchom et al., 2018). Regardless of the source, a multitude of intracellular events are triggered when the DNA in a cell becomes damaged. Cells may, for example, respond to DNA damage by activating DNA repair mechanisms, cell cycle arrest or, in cases of severe DNA damage, apoptosis (Carrassa and Damia, 2017). Such cellular responses to DNA damage are mainly governed by the DNA damage response (DDR), which comprises a complex network of signalling pathways (Carrassa and Damia, 2017). The DDR has many functionalities and, amongst other things, it monitors DNA integrity and repairs DNA damage in order to maintain genomic stability in cells. The DDR also governs DNA replication, cell cycle progression, and apoptosis (Minchom et al., 2018; Nam et al., 2018). When DNA repair in a cell is needed, the DDR activates relevant effector proteins (Minchom et al., 2018). Included in the group of DDR-associated effector proteins are approximately 450 proteins (Nam et al., 2018), out of which the two main regulators for cell cycle checkpoints are ataxia telangiectasia mutated kinase (ATM) and ataxia telangiectasia mutated and rad3-related kinase (ATR) (Sundar et al., 2017). ATM and ATR belong to the enzyme family phosphatidyilinositol-3-OH-kinases (PI3K), and they both play central roles when cells respond to DNA damage (Carrassa and Damia, 2017). In this work, we study the effects of an anti-cancer drug, namely AZD6738, that works by inhibiting ATR activity.

DNA lesions in form of single-strand breaks are a common result of replication stress, and the repair of single-strand DNA breaks is mainly attributed to ATR activity. A drug that inhibits ATR activity consequently inhibits the repair of single-strand DNA breaks post replication stress. Cancer cells are associated with high replication stress and consequently ATR inhibitors have, during the last decade, been explored as anti-cancer agents (Minchom et al., 2018; Carrassa and Damia, 2017; Mei et al., 2019). With the premise that inhibiting DNA damage responses should increase the effect of some other main therapy, DDR inhibitors have been explored as both radiotherapy and chemotherapy treatment intensifiers (Carrassa and Damia, 2017; Mei et al., 2019). Two well-studied ATR inhibitors are AZD6738 and VX-970. AZD6738 is an oral ATR inhibitor, and its anti-tumour potential has been demonstrated in preclinical *vitro* and *in vivo* xenograft studies for various ATM deficient cell lines, including ATM deficient lung cancer, chronic lymphocytic leukemia and metastatic adenocarcinoma of the colon (Checkley et al., 2015; Sundar et al., 2017; Foote et al., 2018). Combination treatments that combine AZD6738 with either radiotherapy or chemotherapy have produced synergistic results in preclinical settings (Sundar et al., 2017), and AZD6738 is currently being evaluated in clinical phase I/II trials (Minchom et al., 2018; Mei et al., 2019). VX-970 is an intravenous ATR inhibitor (Tu et al., 2018) that has demonstrated tumour controlling effects in a phase I clinical trial, both as a monotherapy and in combination with the chemotherapy drug carboplatin (Minchom et al., 2018). A summarising table of clinical trials involving ATR-inhibitors can be found in an article by Mei *et al*. (Mei et al., 2019).

## 2 Model and Method

We use an ABM approach to model monolayer populations of cancer cells, and multi-cellular tumour spheroids that evolve in time and space. The model describes the behaviour of cancer cells using a set of modelling rules. In order to account for differences between monolayer and spheroid scenarios, these rules are adjusted when moving between monolayer and spheroid simulations, as is described throughout Section 2. Taking a minimal parameter approach, we aim to use as few rules and parameters as possible to capture the nature of the regarded systems. We here chose to include model rules and parameters that pertain to the cells’ doubling time and cell cycle (Section 2.2), cell proliferation on the lattice (Section 2.3), the distribution of oxygen and drugs across the lattice (Sections 2.4 and 2.5 respectively) and cellular responses to local oxygen and drug concentrations (Sections 2.4 and 2.6 respectively). In this work, details concerning nutrient distribution and its effect on tumour growth are not included. Instead, under a simplifying modelling assumption, the diffusion of oxygen forms a surrogate for the distribution of nutrients. Differences between monolayer and spheroid simulation modelling rules are pictorially summarised in Section 2.8, and monolayer-calibrated model parameters are listed in Section 2.7.

The *in vitro* and *in vivo* data used in this study are gathered from previous work by Checkley *et al*. (Checkley et al., 2015). In the regarded *in vitro* experiments, populations of LoVo cells were plated and subjected to AZD6738, where population sizes of up to roughly 4000 cells were reported (Checkley et al., 2015). In the *in vivo* experiments, LoVo cells were subcutaneously injected in flanks of female Swiss nude mice in order to produce human tumour xenografts, and AZD6738 treatments started when the tumours had reached a volume of 0.2-0.3 cm^3^ (Checkley et al., 2015). Here, we regard treatment responses in terms two dynamic variables: population cell count or tumour size, and percentage of DNA-damaged (i.e. *γ*H2AX-positive) cells. The *in vitro* and *in vivo* data used in our current study are available in the Supplementary Material (Supplementary Material, S1).

### 2.1 The ABM lattice

In the model, one *agent* corresponds to one cancer cell (in the monolayer simulation) or, due to computational costs, one group of cancer cells (in the spheroid simulation). The behaviour and fate of each agent is governed by a set of rules that incorporate both intracellular and environmental dynamic variables using multiscale modelling techniques (Rejniak and Anderson, 2011). At the start of an *in silico* experiment, one initial agent is placed in the centre of the lattice. This initial agent produces daughter agents and ultimately gives rise to a heterogeneous population of agents. When the population has reached an appropriate size (chosen to match the *in vitro* and *in vivo* data in the monolayer and spheroid simulations respectively), AZD6738 anticancer treatments commence. The ABM lattice is a square lattice, and every lattice point is either empty or occupied by one agent. If a lattice point is empty, it consists of extracellular solution providing nutrients to cells. In the monolayer simulations, the dispersion of any molecules across the lattice is modelled as instantaneous, and thus the extracellular solution is considered to render the entire lattice homogeneous in terms drug and oxygen concentrations at all times. In the spheroid simulations, however, drug and oxygen molecules are modelled as diffusing over the spheroid and the extracellular environment, and consequently the spheroid lattice will be heterogeneous in terms of drug and oxygen concentrations. Oxygen and drug distribution across the lattice are further discussed in Sections 2.4 and 2.5 respectively. Since Checkley *et al*. (Checkley et al., 2015) report monolayer cell population sizes in units of number of cells, and *in vivo* tumour sizes in *cm*^3^, we here choose to measure simulated monolayer and spheroid sizes using cell counts and volumes respectively. The ABM lattices are chosen accordingly, as is described below.

#### Monolayer lattice

Cell populations evolve on a two-dimensional square lattice with 100 × 100 lattice points, where the spacing in both spatial directions, *x*_1_ and *x*_2_, corresponds to one cell diameter.

#### Spheroid lattice

We simulate (only) a central cross section of the spheroid as an, approximately circular, disk of cells living on a two-dimensional square lattice. This lattice is specifically an 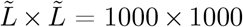 square lattice, with a spacing in both spatial directions 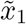 and 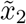 equal to 40*μ*m. The dimensions are chosen in order to allow our ABM to simulate the required physical dimensions, whilst keeping computational costs low. Post simulation time, the twodimensional cross section of cells is extrapolated to represent a three-dimensional spheroid. This disk-to-spheroid extrapolation process is outlined in the Supplementary Material (Supplementary Material, S4).

### 2.2 Cell cycle model

In order to capture the influence of ATR and the ATR inhibitor AZD6738 on the cell cycle, we use a probabilistic, rule-based cell cycle model adapted from previous mathematical (nonagent-based) work by Checkley *et al*. (Checkley et al., 2015). As illustrated in Figure 2, this cell cycle model can be represented as a graph with nodes (cell cycle phases or states) that are connected via various paths (phase/state transitions). A cell can be in an undamaged state (G1, S or G2/M), a replication stress-induced DNA damaged state (D-S) or a dead state, where the cause of cell death is unrepaired replication stress. As ATR is active in the checkpoint in the intra-S phase of the cell cycle, both under undamaged circumstances and in response to DNA damage (Carrassa and Damia, 2017), ATR inhibition will inhibit the cell from progressing to the G2/M state in the mathematical cell cycle model. A cell can take different paths through the cell cycle graph, and every time that paths fork, random number generation determines which path will be taken. Every cell commences its life in the G1 state, but thereafter a cell can enter either the S state or the damaged S (D-S) state. The probability that a cell enters the D-S state is denoted Π_*D–S*_ and is calibrated by *in vitro* data (Checkley et al., 2015). If a cell enters the D-S state, it has a chance to repair itself and enter the S state. If there is no drug in the system, this repair is always achieved, however the repair path is inhibited by the presence of the drug AZD6738. The higher the drug-concentration, the more unlikely it is that a cell in the D-S state will successfully repair itself to the S state. If a cell in the D-S state fails to repair, it is sentenced to die. Whether a cell in state D-S repairs or dies is decided by comparing a random number, generated from a uniform distribution, to the cell’s survival probability, which is influenced by the local drug concentration 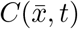, as is described in detail in Section 2.6. A cell that has successfully reached the S state continues to the G2/M state, after which it duplicates and starts over in the G1 state again, ready to perform another cell cycle.

**Figure 2:**
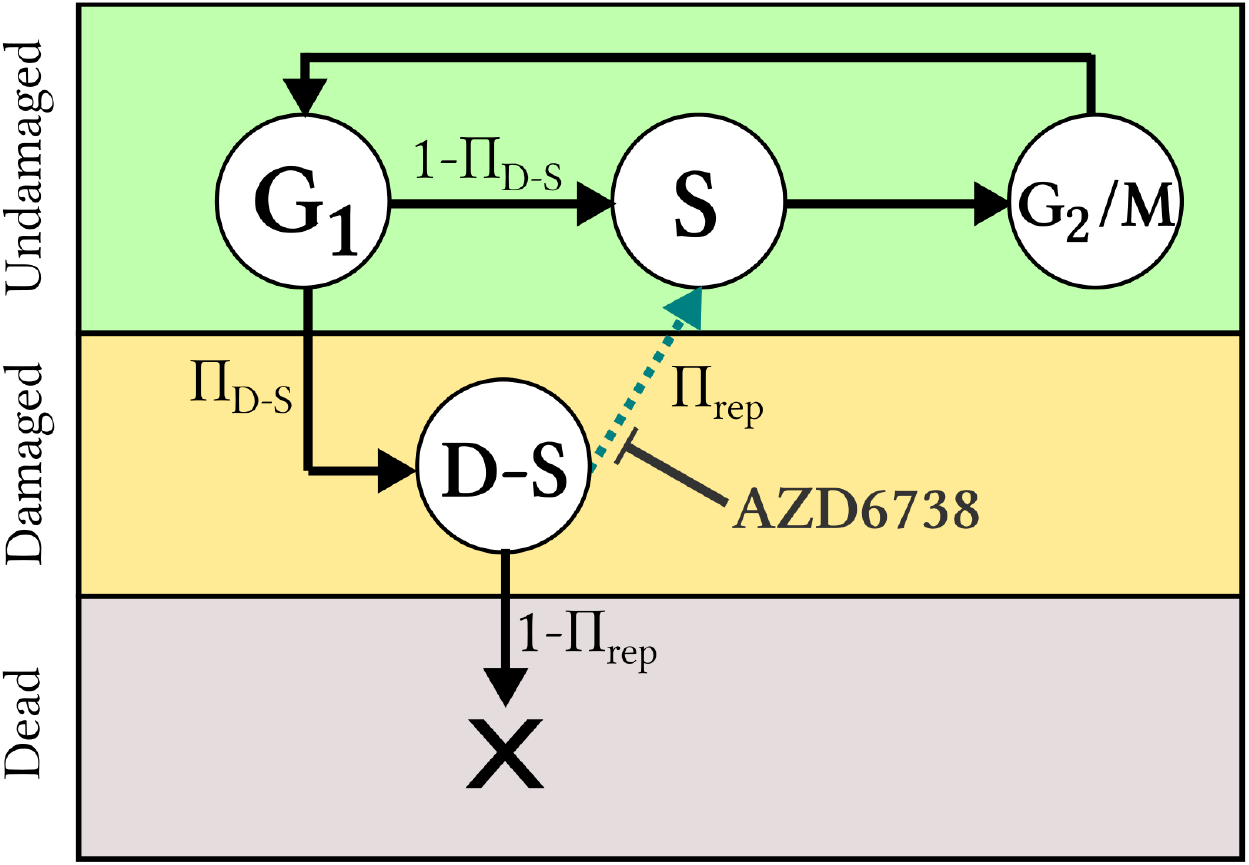
Cell cycle model: An agent, *i.e*., a cell (in the monolayer simulation) or a group of cells (in the spheroid simulation), progresses through various states of the cell cycle, where the states correspond to cell cycle phases and are shown as nodes in the graph. Viable (undamaged or damaged) states are shown in circles, whilst the dead state is shown as a cross. Paths illustrate transitions between states, and symbols next to the paths denote the probabilities that the corresponding paths will be taken. The dashed path can be inhibited by an ATR inhibiting drug, such as AZD6738.

In order to allow for asynchronous populations, each agent *i* on the lattice is assigned an individual doubling time *τ_i_*, where *τ_i_* is a random number generated from a normal distribution with mean value *μ* and standard deviation *σ*. Each agent is attributed an individual cell cycle clock, that determines when the agent should progress to a subsequent state in the cell cycle model. Progression to a subsequent cell cycle state occurs once an agent has spent a certain fraction of its doubling time in its current state. The fraction of the doubling time spent in the G1, S (including D-S) and G2/M states are respectively denoted Θ_*G*1_, Θ_*S*_ and Θ_*G*2/*M*_, where these values are approximate and chosen from literature to match values for typical human cells with a rapid doubling time of 24 hours so that Θ_*G*1_ = 11/24, Θ_*S*_ = 8/24 and Θ_*G*2/M_ = 5/24 (Cooper and Hausman, 2007). The fraction of an agent’s doubling-time spent in the D-S state, Θ_*D–S*_, is on the other hand fitted by *in vitro* data produced by Checkley *et al*. (Checkley et al., 2015), as outlined in the Supplementary Material (Supplementary Material, S2). Monolayer and spheroid cell cycle modelling rules are described below.

#### Monolayer cell cycle model rules

One agent corresponds to one cancer cell that is assigned an individual doubling time *τ_i_*. The cell cycle path taken by cell *i* is governed by drug concentrations and random number generations specific to that cell.

#### Spheroid cell cycle model rules

One agent comprises a group of identical cancer cells. Each agent is assigned an individual doubling time, *τ_i_*, and thus all cells belonging to agent *i* progress simultaneously and uniformly through the cell cycle model. Random number generations specific to agent i determine which path the agent takes through the cell cycle.

### 2.3 Cell proliferation

When an agent has completed the mitosis state in the cell cycle model a daughter agent is produced. Each daughter agent is placed on a random lattice point in the (approximately circular) neighbourhood of its parental agent. To accomplish circular-like growth, the model stochastically alternates between placing daughter agents on Moore and von Neumann neighbourhoods of parental agents, as is pictorially described in the Supplementary Material (Supplementary Material, S3). A daughter agent is allowed to be placed on, up to, a *ν*th order neighbourhood of its parental agent, but lower order neighbourhoods (i.e. neighbourhoods closer to the parent) are prioritised and populated first. Modelling rules concerning monolayer and spheroid cell proliferation are outlined below.

#### Monolayer proliferation rules

In the experimental monolayer *in vitro* setup, there is no spatial constraint or nutrient deficiency that is inhibiting cell division within the time-course of the experiment. Consequently cells are allowed to divide freely in the monolayer model and we set *ν* to be equal to infinity (with the restriction that agents can not be placed outside the lattice in the *in silico* implementation). Although this non-local placement of daughter cells neglects physics, we are not considering spatial heterogeneity in the monolayer simulation and therefore cell location does not affect the evaluated simulation results.

#### Spheroid proliferation rules

*In vivo* tumours and *in vitro* spheroids typically consist of a core with non-proliferating cells and a shell of proliferating cells. To accommodate for this, a daughter agent (representing a group of daughter cells) is allowed to be placed on up to a third order (approximately circular) neighbourhood of its parental agent, so that 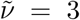, in accordance with previous mathematical models (Powathil et al., 2012b). For the spheroid simulation regarded in our current study, 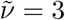 matches the experimental *in vivo* data. However, for other experiments, the value of 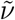 should be adjusted to fit the specific cell-line and modelling scenario at hand. When an agent is in the G1 phase of the cell cycle, it scans its environment to see if it has enough resources, in terms of space and nutrients, to commence the process of producing a daughter cell. If not, the cell enters the quiescent phase (Alarcon et al., 2004). Thus in the model, when an agent is in the G1 phase, it continues to progress through the cell cycle model, provided that some free space is available on the lattice within its 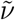th neighbourhood. If this is not the case, the agent exits the cell cycle to enter a quiescent state G0. Should neighbourhood space be made available again, here as a result of anti-cancer targeting, quiescent agents may re-enter the cell cycle.

### 2.4 Oxygen distribution and influence on cells

Tumour growth and treatment responses are highly influenced by intratumoural oxygen levels (Hu et al., 2010; Liapis et al., 2015; Peeters et al., 2015) and severely hypoxic (cancer) cells may proliferate slower than well-oxygenated cells (Alarcon et al., 2004).

#### Monolayer oxygen distribution and responses

In the mathematical monolayer model, all cells are assumed to be well-oxygenated in accordance with the experimental *in vitro* setup performed by Checkley *et al*. (Checkley et al., 2015). Consequently, neither oxygen dynamics nor cellular responses to oxygen levels are incorporated in the monolayer model.

#### Spheroid oxygen distribution and responses

Within solid tumours, oxygen concentrations typically vary and hypoxic regions are common tumour features (Peeters et al., 2015; Hamis et al., 2020a; Sun et al., 2012). Oxygen gradients are also observed in *in vitro* tumour spheroids (Voissiere et al., 2017), thus oxygen dynamics across the lattice are here described using a mechanistic diffusion equation, where the oxygen concentration in location 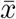 at time *t* is denoted by 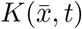 where

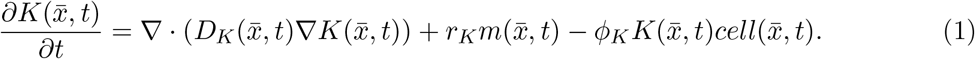

The first term in Equation 1 describes oxygen diffusion across the lattice, the second term is an oxygen supply term and the third term describes oxygen uptake by cells. Accordingly, 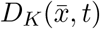 is the oxygen diffusion coefficient, and *r_K_* and *ϕ_K_* are supply and consumption coefficients respectively. The diffusion coefficient for oxygen is known from literature to be 2.5 × 10^-5^ cm^2^s^-1^ (Powathil et al., 2012b). Assuming that oxygen diffuses slower inside the spheroid than outside the spheroid, the oxygen diffusion coefficient is divided by a factor 1.5 if there is a cell in location 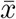 at time *t* (Powathil et al., 2012b). The binary factor 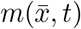 is 1 if the regarded location 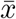 is outside the spheroid boundary at time *t* and 0 otherwise, *i.e*., 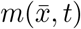 is 1 if the regarded lattice point is not occupied by an agent nor completely surrounded by agents, thus oxygen is here modelled as supplied from ‘outside the boundary of the spheroid’. Similarly, the binary factor 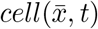 is 1 if there is a viable (non-dead) cell in location 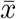 at time *t*, and 0 otherwise (Powathil et al., 2012b). Equation 1 is coupled with no-flux boundary conditions, thus the total amount of oxygen in the system will fluctuate over time (Powathil et al., 2012a). A scaled oxygen variable 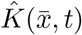 is introduced in order to express oxygenation levels in percentages (%) between 0% and 100%. This scaled oxygen value is computed at every unique time step *t_u_* by

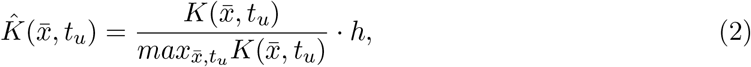

where 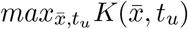 denotes the maximum occurring 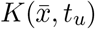-value at the time point *t_u_* and *h* is a scaling factor that is included in order to classify cells with oxygen levels of 10%, or less, as hypoxic (Powathil et al., 2012b; Hamis et al., 2020a). Low cellular oxygen levels have been shown to delay cell cycle progression by inducing arrest in, particularly, the G1 phase of the cell cycle (Alarcon et al., 2004). Consequently, in our model, hypoxic cells display arrest (i.e. delay) in the G1 phase of the cell cycle. In mechanistic Tyson-Novak type cell cycle models (Tyson and Novak, 2001; Novak and Tyson, 2003, 2004), the cell cycle is governed by a system of ordinary differential equations (ODEs) in which the G1 phase can be inherently elongated under hypoxic conditions by incorporating hypoxia-induced factors into the ODEs (Powathil et al., 2012b). In the mathematical model discussed in this paper, however, we use agent-attributed clocks to model cell cycle progression and thus, in order to achieve a longer G1-phase under hypoxic conditions, we introduce a G1 delay factor (G1DF) (Hamis et al., 2020a) where

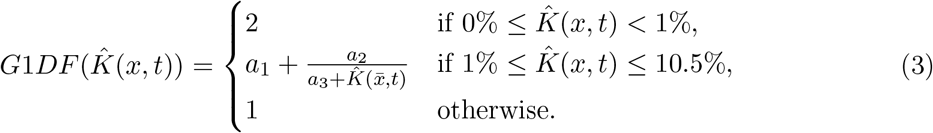

The G1DF is an approximation for how much the G1 phase is expanded in time as a function of cellular oxygen concentration. It is matched to fit data points extracted from a previous mathematical study by Alarcon *et al*. (Alarcon et al., 2004), in which a Tyson-Novak cell cycle model is extended to incorporate the action of p27, a protein that is up-regulated under hypoxia and delays cell cycle progression. Data-fitting yields the parameter values *a*_1_ = 0.9209, *a*_2_ = 0.8200, *a*_3_ = –0.2389 (Hamis et al., 2020a). Thus the fraction of an agent’s doubling time spent in the G1 state is 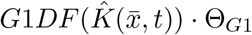, where 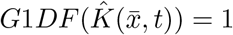 for normoxic cells.

### 2.5 Drug distribution across the lattice

Drug distribution significantly varies between monolayer and spheroid settings. In the regarded monolayer setup, the drug concentration can be regarded as homogeneous, whilst heterogeneous drug concentrations must be accounted for when simulating drug distribution across the spheroid. Drug uptake and receptor dynamics is omitted in the model, the drug response of an agent is instead influenced by the drug concentration in the lattice point that it occupies.

#### Monolayer drug distribution

In the *in vitro* experiments performed by Checkley *et al*. (Checkley et al., 2015), plated cell populations of roughly 1000 cells were treated with AZD6738 in the solvent dimethylsulfoxide (DMSO). In the mathematical model, we approximate the drug distribution across the lattice to be instantaneous (occurring at treatment time *T*_0_) and homo-geneous. We furthermore assume that the drug has a half-life time that exceeds the time course of the experiment, and note that there is no other drug elimination from the *in vitro* system. In our mathematical model, this is equivalent to there being no drug decay or elimination, hence the drug concentration 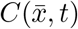, in location 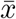 at time *t* is simply given by

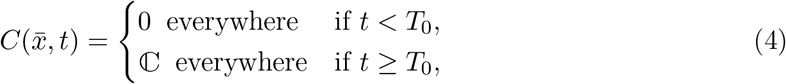

where 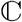 denotes the applied drug concentration (in units of molarity).

#### Spheroid drug distribution

The spheroid scenarios simulated in this study are compared to the in vivo experiments performed by Checkley *et al*. (Checkley et al., 2015), in which the drug AZD6738, or vehicle in the control case, were administered via oral gavage once per day to female Swiss nude mice. Therefore we include dynamic drug delivery and drug decay in our spheroid simulations. In the mathematical spheroid model, we consider the drug to diffuse through the spheroid from its surrounding, creating a drug gradient within the spheroid. This drug dynamics is modelled using a partial differential equation (PDE), where the concentration of AZD6738 at location 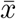 at time *t* is denoted by 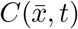 such that

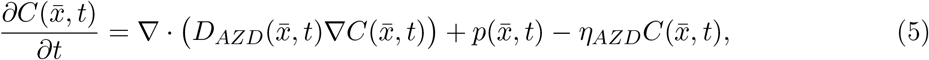

where *D_AZD_* is the diffusion coefficient of the drug AZD6738, and the supply coefficient 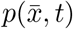 is greater than zero at drug administration times only for lattice points outside the tumour. Assuming first order kinetics for drug elimination, the drug decay constant *η_AZD_* is matched to the reported half-life time of 6 hours for AZD6738 *in vivo* (Vendetti et al., 2015). Note that the drug decay term here represents all drug elimination from the system, both metabolic and that caused by excretion.

The diffusion rate of a drug is predominantly affected by the molecular size of the drug. More specifically, the diffusion coefficient of a drug is inversely proportional to the square root of the molecular weight of the drug, so that large molecules diffuse more slowly than do small molecules (Dale and Rang, 2007). Using this assumption, the drug diffusion coefficient *D_AZD_* is set in relation to the oxygen diffusion coefficient *D*_0_2__, as is done in previous mathematical studies (Powathil et al., 2012b). Thus the relationship between the diffusion coefficients corresponds to the square of the inverse relationship between the molecular weights, such that

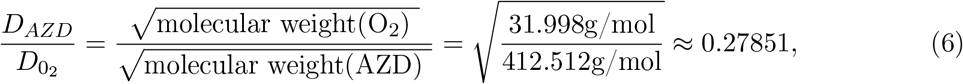

where the molecular weights are collected from the PubChem database (Kim et al., 2019). Details regarding pharmacokinetics are outside the scope of this study, bioavailability is instead calibrated using extreme case drug scenarios, as described in the Supplementary Material (Supplementary Material, S2).

### 2.6 Drug responses

AZD6738 inhibits DNA repair from the D-S state to the S state in the cell cycle model, as illustrated in Figure 2, and, in our model, maximal drug effect corresponds to complete repair inhibition. The drug effect is modelled using an agent-based adaptation of the sigmoid Emax model (Holford, 2017), a note on the choice of drug model is included in the Supplementary Material (Supplementary Material, S6). In the ABM-adapted sigmoidal Emax model used here, the drug effect on a cell in position 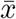 at time *t* is given by

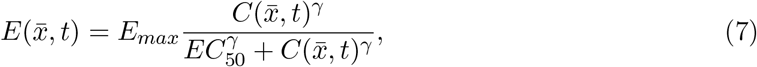

where the drug concentration in lattice point 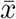 at time *t* is given by 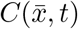. *E_max_* denotes the maximal drug effect, here corresponding to complete repair inhibition (*E_max_* = 1), *EC*_50_ denotes the drug concentration required to achieve half of the maximal drug effect (0.5 · *E_max_*) and *γ* is the Hill-exponent (Holford, 2017). *EC*_50_ and *γ* are fitted from the *in vitro* data, as outlined in the Supplementary Material (Supplementary Material, S2). When an agent is scheduled to progress from the D-S state in the cell cycle, it has a probability Π_*rep*_ ∈ [0,1] to repair, where Π_*rep*_ is determined by the local drug concentration so that

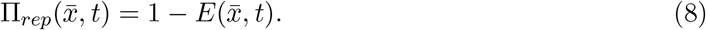

Note that in the absence of drugs, the repair probability is 1. When a cell dies, it is transformed into a membrane-enclosed ‘cell-corpse’ (Dale and Rang, 2007). In an *in vivo* setting, this cellular debris is digested by macrophages but in an *in vitro* setting such ‘cell-corpses’ may linger on the lattice during the course of the experiment. Post the lethal event (i.e. the D-S to S repair failure) a cell is declared ‘dead’ in the model after a time *T_L→D_* has passed (where *L* stands for ‘lethal event’ and *D* stands for ‘death’). The parameter *T_L→D_* is calibrated by *in vitro* experiments. The differences between modelling rules for monolayer and spheroid drug responses are described below.

#### Monolayer drug responses

After failure to repair from the D-S state, a cell (*i.e*., an agent) is considered to be dead after a time *T_L→D_* has passed. However, a dead cell is never physically removed from the lattice.

#### Spheroid drug responses

In order to simulate *in vivo*-like removal of dead cancer cells, an agent (*i.e*., a group of cells) is declared to be dead and is removed from the lattice after an amount of time *T_L→D_* post the lethal event (failure to repair).

### 2.7 Parameters

The parameters used in the mathematical model are calibrated by monolayer *in vitro* data, this calibration process is described in the Supplementary Material (Supplementary Material, S2). In the context of quantitative pharmacology, knowledge about a model’s robustness is crucial (Visser et al., 2014), therefore we have provided results from the uncertainty and sensitivity analysis in the Supplementary Material (Supplementary Material, S9). We performed three different uncertainty and sensitivity analyses techniques, suitable for stochastic agent-based models, namely (*i*) consistency analysis, (*ii*) robustness analysis and (*iii*) Latin hypercube analysis (Hamis et al., 2020b; Alden et al., 2013). Detailed descriptions on how to perform and interpret these techniques are available in an introductory uncertainty and sensitivity analyses review (Hamis et al., 2020b). In accordance with the performed consistency analysis, we run 100 simulations per *in silico* experiment in order to formulate results (in terms of mean values and standard deviations) that mitigate uncertainty originating from intrinsic model stochasticity.

### 2.8 Differences between monolayer and spheroid modelling rules

Modelling rules are adjusted when moving between monolayer and spheroid simulations. Differences between monolayer and spheroid rules are pictorially summarised in Figure 3. A note on the simplifying modelling assumptions that we have used in this study is provided in the Supplementary Material (Supplementary Material, S5).

**Figure 3:**
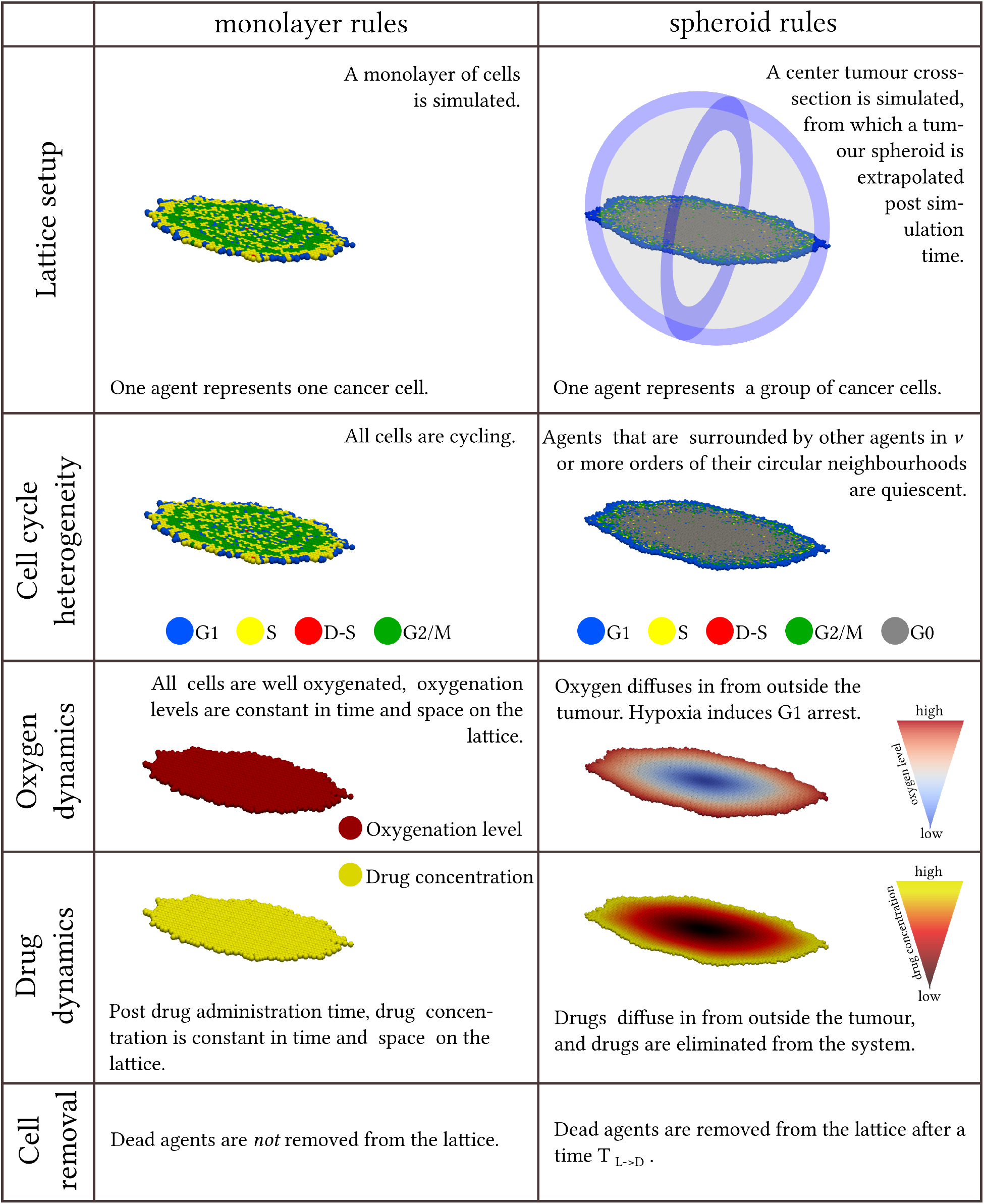
A summary of the differences between the monolayer and spheroid modelling rules used in the mathematical framework.

### 2.9 Implementation

The mathematical model is implemented in an in-house C++ (Stroustrup, 1995) framework, in which PDEs are solved using explicit finite difference methods. Simulation cell-maps are visualised using ParaView (Ayachit and Utkarsh, 2015) and data analysis, as well as uncertainty and sensitivity analyses, are performed in MATLAB (MATLAB, 2019).

## 3 Results

The mathematical framework is first calibrated by *in vitro* monolayer data produced by Checkley *et al*. (Checkley et al., 2015). Thereafter, spheroids subjected to dynamic drug delivery and the removal of dead cells are simulated. Spheroid simulations are then compared to *in vivo* treatment responses in human tumour xenografts. Two model outputs are considered in the *in silico* simulations: the fraction of DNA damaged cells in the system and the size of the cancer cell population or tumour spheroid over time. Note that, in the model, a cell is classified as DNA-damaged if it is in the D-S state of the cell cycle depicted in Figure 2. In the experimental setup, DNA damaged cells are labeled as *γ*-H2AX positive (Checkley et al., 2015).

### 3.1 Simulating monolayer experiments

In the *in vitro* experiments, populations of LoVo (human colon carcinoma) cells were exposed to the ATR inhibiting drug AZD6738 (Checkley et al., 2015). Figure 4 shows monolayer simulation results, specifically the percentage of DNA damaged (*γ*H2AX-positive) cells over time (Left) and the total cell count over time (Right). In the simulations, AZD6738 drugs are given at 0 hours, when the cell population has reached a size of approximately 1000 cells. Simulated response curves for six different drug concentrations, including the zero-drug concentration control case, are shown. Also shown in Figure 4 are *in vitro* data and results from the mathematical compartment-ODE model presented by Checkley *et al*. (Checkley et al., 2015) describing the same experimental scenario. Using a minimal-parameter modelling approach, the mathematical framework is calibrated to fit *in vitro* data points without introducing any variable model parameters. This calibration process is described in the Supplementary Material (Supplementary Material, S2).

**Figure 4:**
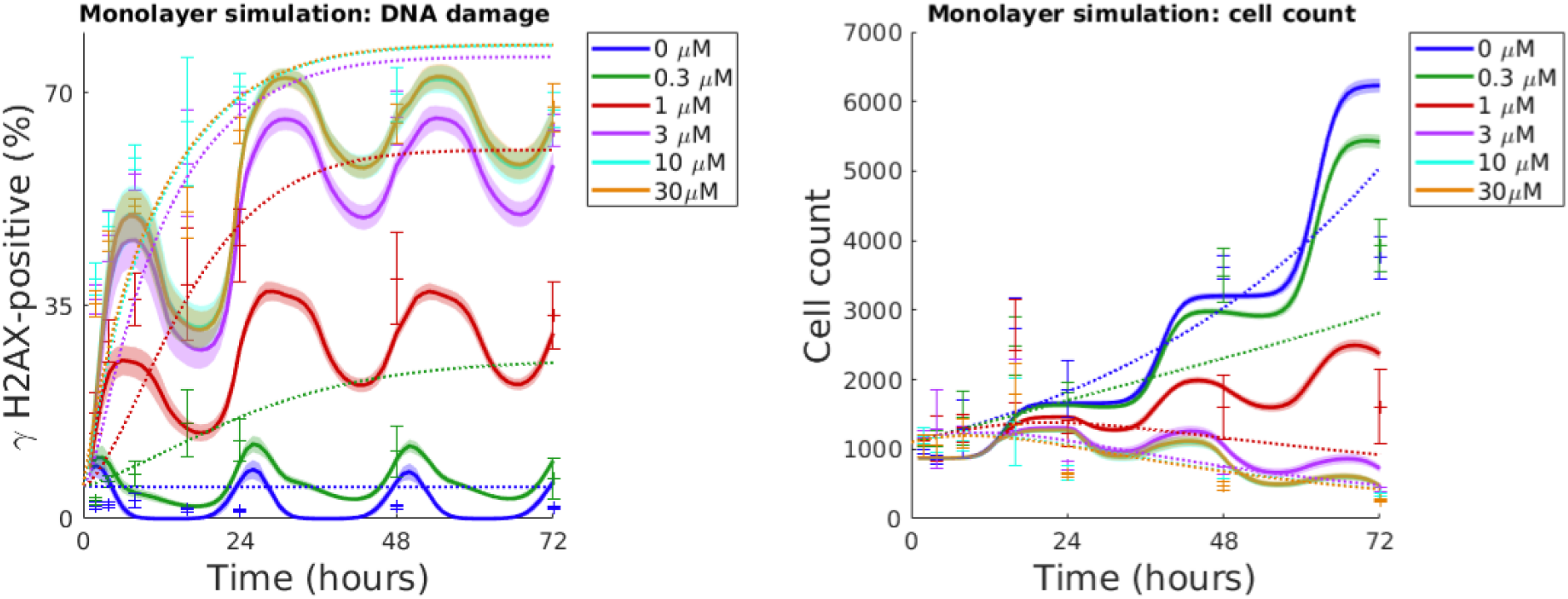
Simulated monolayer drug response curves are compared to *in vitro* data and mathematical results from a previously published compartment-ODE model by Checkley *et al*. (Checkley et al., 2015). LoVo cells are exposed to drug (AZD6738) at 0 hours. Left: The percentage of *γ*H2AX-positive (DNA-damaged) cells in the system over time. Right: Cell count over time. ABM simulated mean values and standard deviations for 100 *in silico* runs are shown with solid lines and shaded ribbons respectively. *In vitro* data in form of mean values and standard deviations are demonstrated with center points and error bars (Checkley et al., 2015). Compartment-ODE model results are represented by dotted lines.

Our results demonstrate that, post *in vitro* monolayer parameterisation, our mathematical framework is able to capture the qualitative nature of *in vitro* monolayer LoVo cell population growth and drug (AZD6738) responses. The model qualitatively reproduces the asymptotic fraction of DNA damaged cells in the system but fails to match early data points (Figure ?? Left). The sensitivity analysis (Supplementary Material, S9) demonstrates that the treatment timing (in relation to the overall cell cycle phase composition of the cancer cell population) notably influences treatment responses in terms of percentage of *γ*H2AX-positive cells. The model parameter calibration process selects for a strong cell cycle synchronisation amongst cancer cells, determined by the model parameter *σ* (Table 1). This strong synchronisation gives rise to oscillatory cell cycle state compositions, as can be seen in Figure 2 where cell cycle specific cell counts are plotted over time in response to different drug doses. This strong synchronisation also yields the step-wise growth curves seen in Figure 4 (Right). The experimental error bars in this figure, and the numerical cell count data available in Table S1 (Supplementary Material, S1), demonstrate that the doubling time of the cell population drastically decreased towards the end of the *in vitro* experiment and, consequently, our agent-based model was not able to replicate cell count data at 72 hours as the modelling rules and parameters are not updated over time.

**Table 1:** *In vitro* calibrated parameters. *μ* and *σ* respectively denote the mean value and the standard deviation of the normal distribution from which an agent’s cell cycle length is picked. Π_*D–S*_ and *θ_D–S_* denote the probability that an agent enters the D-S state, and the fraction of its cell length spent in the D-S state. EC_50_ and *γ* denote the half maximal drug concentration and the Hill-exponent in the drug response equation (5). *T_L→D_* denotes the time it takes for an agent to die post DNA damage repair failure, and *τ_i_* denotes the cell cycle length of agent *i*.

| Section | Parameter | Calibrated Value |
| --- | --- | --- |
| 2.2 | $\mu, \sigma$ | 24 h, 0.5 h |
| | $\Pi_{D-S}, \theta_{D-S}$ | 0.75, 0.03 |
| 2.6 | $EC_{50}, \gamma$ | 1 μM, 2 |
| | $T_{L \rightarrow D}$ | $\tau_i$ |

The ABM model and Checkley *et al*.’s (Checkley et al., 2015) compartment-ODE model are also compared to each other and *in vitro* data in residual plots available in the Supplementary Material (Supplementary Material, S7). In an effort to quantify how well the two mathematical models match the data, the Root Mean Square Errors (RMSEs) are computed between *N* simulation mean values and data mean values so that 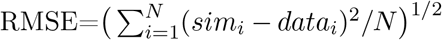. When comparing DNA damage simulation results to data, the ABM model yields an RMSE of 11.6 percent units, whilst the compartment-ODE model yields an RMSE of 14.6 percent units. When comparing cell count simulation results to data, however, the ABM RMSE is 644 cells whilst the compartment-ODE RMSE is 495 cells.

### 3.2 Simulating spheroid experiments

Post *in vitro* monolayer calibration, the mathematical framework is used to simulate spheroid experiments, that are compared to the *in vivo* experiments performed by Checkley *et al*. (Checkley et al., 2015) in which LoVo xenografts, that are injected in mice flanks, are treated with AZD6738 once daily for 14 days. The results in Figure 6 show AZD6738 drug responses in terms of the percentage of DNA damaged (γH2AX-positive) cells (Left) and spheroid/tumour volume (Right) over time. Simulated response curves to three different drug doses (0, 25 and 50 mg/kg) and *in vivo* data are provided in Figure 6.

**Figure 5:**
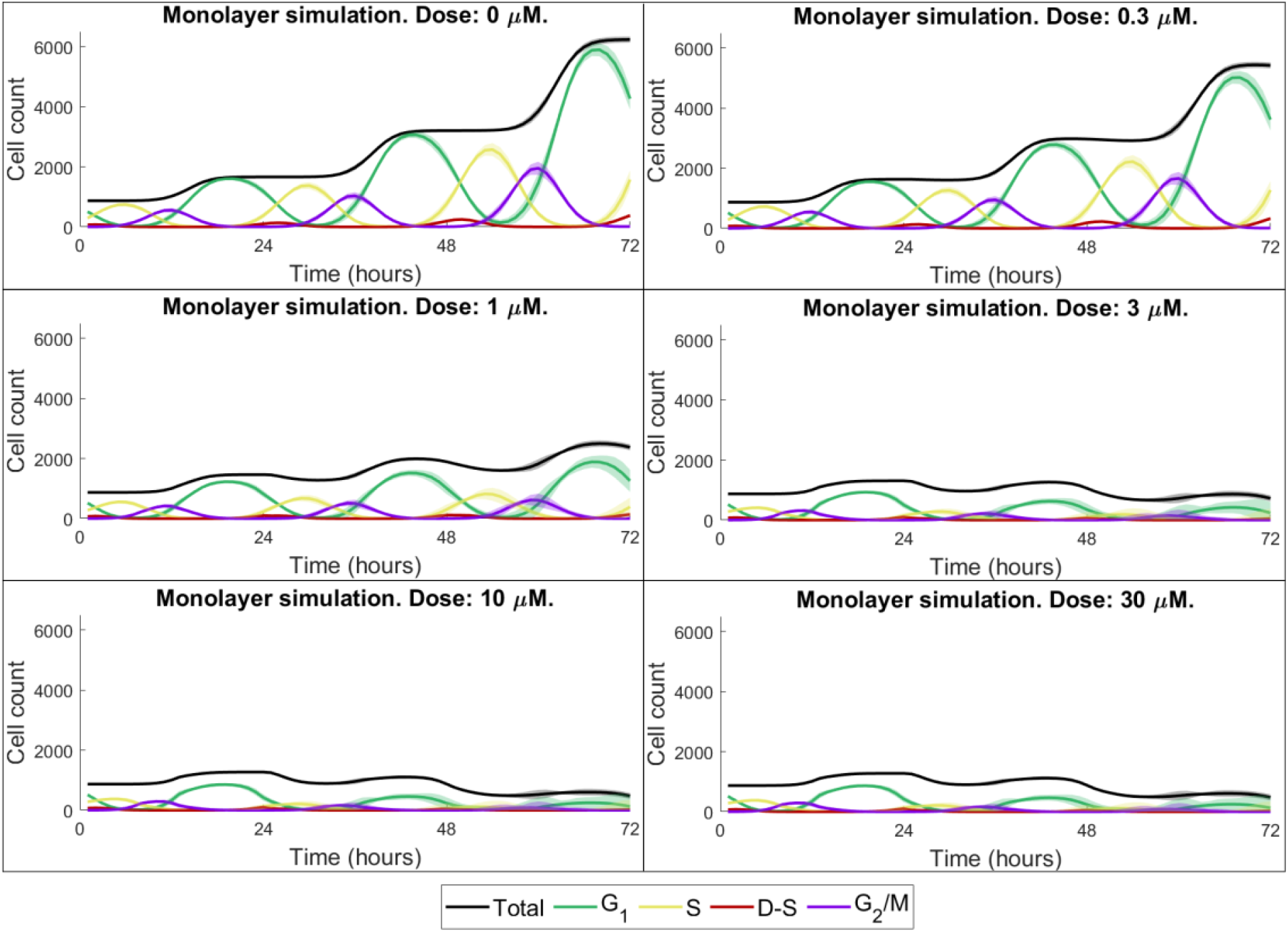
Cell cycle state specific monolayer cell counts. Each subplot shows the number of cells in the G_1_, S, D-S, G_2_/M state, as well as the total cell count, for a specific drug dose. Mean values and standard deviations for 100 *in silico* runs are shown with solid lines and shaded ribbons respectively.

**Figure 6:**
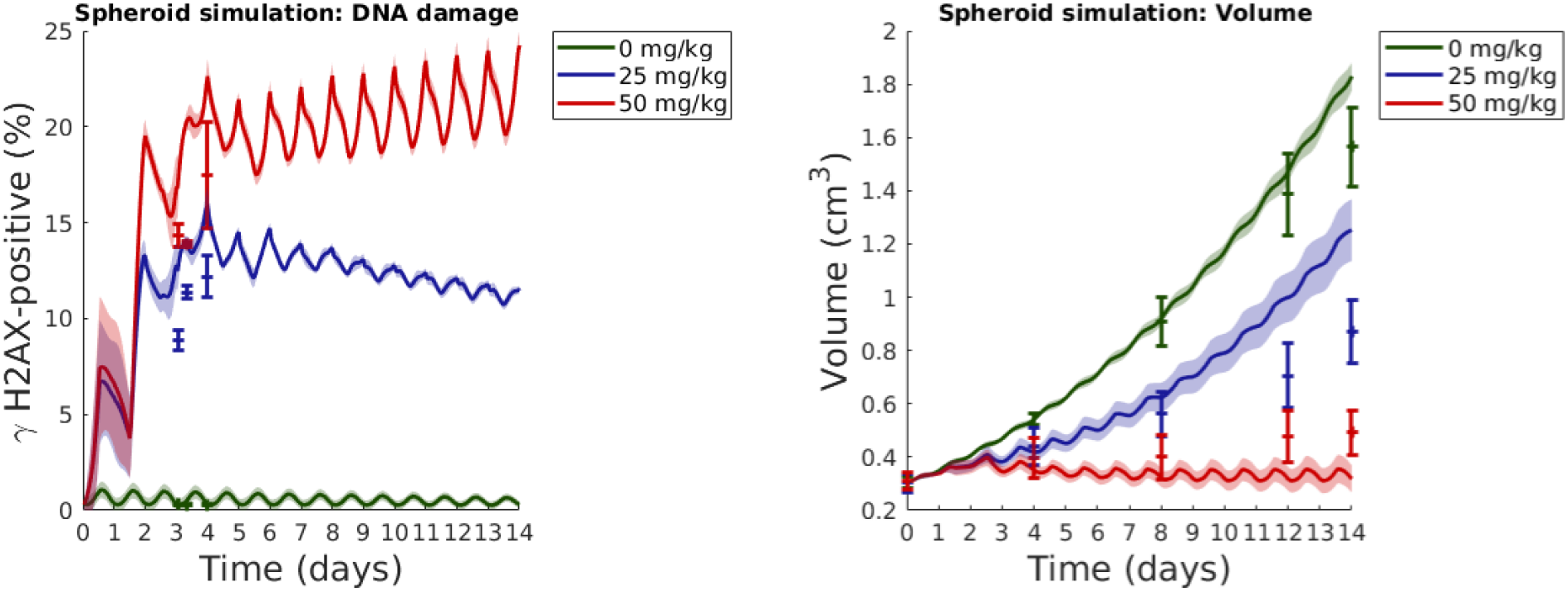
Simulated spheroid drug response curves are compared to *in vivo* xenograft data. In both the spheroid simulation and the *in vivo* experiment, LoVo xenografts are exposed to drug (AZD6738) once daily for 14 days. Left: The percentage of γH2AX-positive (DNA-damaged) cells in the spheroid/xenograft over time. Right: Spheroid/xenograft volume over time. Simulated (spheroid) mean values and standard deviations for 100 *in silico* runs are shown with solid lines and shaded ribbons respectively. *In vivo* data in form of mean values and standard errors are demonstrated with center points and error bars (Checkley et al., 2015).

Figure 6 (Right) demonstrates that our simulated spheroid results qualitatively agrees with the *in vivo* results reported by Checkley *et al*. (Checkley et al., 2015) for approximately 12 days post tumour injection for control case tumours, and for approximately 8 days post tumour injection for tumours subjected to drugs. This can be explained by the fact that the behaviour of the agents in our current model does not change over time, when in fact tumours are highly adaptable and responsive to external pressures. It follows that details pertaining to tumour growth and drug sensitivity may vary over time, and in future work the agent-based model used in this study can be updated to incorporate variable modelling rules and parameters. Residual plots, comparing ABM simulation results to *in vivo* data (Checkley et al., 2015) are available in the Supplementary Material (Supplementary Material, S7) for DNA damage and spheroid/tumour volume evaluations. The RMSE for DNA results is 3.57 percent units. For the spheroid/tumour volume simulation results, the RMSE is 0.038 cm^3^ up to and including 8 days, but 0.16 cm^3^ for the full 14 day simulations.

## 4 Discussion

Simulation results obtained in this study were compared to *in vitro* and *in vivo* data and, furthermore, to previous mathematical modelling results produced by Checkley *et al*. (Checkley et al., 2015). In their study, Checkley *et al*. (Checkley et al., 2015) modelled tumour responses to AZD6738 using coupled ordinary differential equations, where a pharmacokinetic/pharmacodynamic (PK/PD) model of tumour growth was integrated with a mechanistic cell cycle model. Their model is predictive of *in vivo* xenograft studies and is being used to quantitatively predict dose and scheduling responses in a clinical Phase I trial design (Checkley et al., 2015). Our modelling results qualitatively agree with those produced by Checkley *et al*. (Checkley et al., 2015), although two different modelling approaches have been taken: Checkley *et al*. (Checkley et al., 2015) regard the tumour as one entity with different compartments whilst we here use a bottom-up modelling approach and regard the tumour as consisting of multiple, distinct agents. Since AZD6738 specifically targets cells that are in the damaged S cell cycle state, we included cell cycle phase resolution into the ABM. Although this modelling approach makes parameter calibration more difficult (compared to more phenomenological models in which the drug acts on all cells), it provides an opportunity to study details about the biological system *in silico* that are not easily observable *in vitro* or *in vivo*. Cell cycle phase details will furthermore be of importance in future work in which the model will be extended to include combination treatments, as many anti-cancer treatments are cell cycle phase specific (Mills et al., 2018).

Moving drug-response investigations from *in vitro* to *in vivo* settings is a key step involved in the process of moving a drug from *bench-to-bedside*. However, *in vivo* data are often sparse, as gathering *in vivo* data is associated with practical, financial and ethical constraints. Plentiful and adaptable *in silico* data are, on the other hand, easy to produce, and can thus be used as epistemic complements to sparse *in vivo* data. Well-formulated *in silico* tools can be extended to investigate various dose-schedule scenarios in order to guide *in vitro* and *in vivo* experiments. Such *in silico* experiments may provide a testbed for simulating various mono and combination therapies. In this study we aimed to capture treatment responses in tumour spheroids (with dynamic drug delivery and the removal of dead cells) using monolayer data and modelling rules that are based on chosen “fundamental” principles that describe how cancer cells in a system behave. Although our spheroid simulations were able to qualitatively mimic the dynamics of *in vivo* xenografts at early time-points (up to 8 days) post tumour injection, the model did not match data at later time points. This is to be expected, as the effects of certain biological processes that are present *in vivo*, but not *in vitro* (*e.g*., angiogenesis), do not impact the tumour volume instantly after tumour injection. Thus the spheroid model can, in future work, be extended to more accurately simulate *in vivo* scenarios. For example, stromal tumours cells, angiogenesis and metastasis can be included in the model. Moreover, further heterogeneity amongst cancer cells can be incorporated, pertaining to *e.g*. drug resistance-related variables. In order to account for mechanical aspects of tumour growth, the approximated cancer cell population/tumour growth model, which allows for daughter agents being placed on non-adjacent lattice point of the parental agent, can be updated to a more realistic proliferation model. Pharmacokinetic details, drug uptake and receptor dynamics can be included in order to make the drug model more detailed. In order to realistically simulate *in vivo* tumours, effects of the host’s immune system can furthermore be incorporated in the model. The observation that anti-cancer drug responses vary between *in vitro* monolayer, *in vitro* spheroid and *in vivo* models has also been addressed in a mathematical study by Wallace *et al*., who used ODE models to simulate neuroblastoma treated with 15-Deoxy-*PGJ*_2_ in monolayers and spheroids (Wallace et al., 2016). Similar to our modelling approach (Figure 1), the authors used *in vitro* data to calibrate a monolayer model, and thereafter extended the model to incorporate spheroid features in order to simulate spheroid dynamics.

The ABM considered in this study is an extension of a mathematical model that has previously been used to study tumour growth and treatment responses to chemotherapy, radiotherapy, hyperthermia and hypoxia-activated prodrugs (Powathil et al., 2012b; Hamis et al., 2020a; Powathil et al., 2015; Hamis et al., 2018, 2019; Bruningk et al., 2018). In recent years, several ABMs have been developed for the purpose of describing various aspects of cancer dynamics (Metzcar et al., 2019), and it should be noted that the modelling approach proposed in Figure 1 is not conceptually limited to usage with the ABM described in this study. The choice of ABM should be influenced by the research question at hand, the desired level of model details and the available data. Examples of data-driven ABMs are available in a recent review article by Chamseddine and Rejniak (Chamseddine and Rejniak, 2020) discussing hybrid models, and hybrid modelling techniques, used in the field of mathematical oncology today.

Data-driven modeling, exploitation of existing data and proof-of-concept studies are important steps involved in current and future procedure for enabling mathematical modeling in systems medicine, as argued in a report by Wolkenhauer *et al*. (Wolkenhauer et al., 2014). A pipeline for predicting therapy outcomes using data-driven mathematical modelling is proposed by Brady and Enderling (Brady and Enderling, 2019) in a recent publication. Despite the fact that mathematical modelling is becoming increasingly popular in the pharmaceutical industry, there are not that many ABMs present in the pharmaceutical scene (Cosgrove et al., 2015). We argue that this is a missed opportunity in the context of oncology, as ABMs naturally capture the heterogeneous nature of tumours, which is known to complicate treatments. As multiscale ABMs organically enable the integration of data across various scales in time and space, it follows that they are useful to the interdisciplinary team that wishes to combine data and knowledge from its team members. Following interdisciplinary collaborations between clinicians, biologists and mathematicians, mathematical modelling may be used to enable *in silico* informed drug development.

## Declarations

### Availability of data and material

All *in vitro* and *in vivo* data used in this study are gathered from Checkley *et al*. (Checkley et al., 2015) and are listed in the Supplementary Material (Supplementary Material, S1).

### Code availability (software application or custom code)

Project code is available on the code hosting platform GitHub at https://github.com/SJHamis/DDRinhibitors.

