## Supplementary Material S1-S9. for "Targeting cellular DNA damage responses in cancer: An *in vitro*-calibrated agent-based model simulating monolayer and spheroid treatment responses to ATR-inhibiting drugs"

S1. Experimental data

S2. Model parameterisation

S3. Proliferation of agents on the lattice

S4. Cross-section to tumour spheroid extrapolation

S5. A note on simplifying modelling assumptions

S6. A note on the choice of drug model

S7. Residual plots

S8. Spheroid waterfall plot

S9. Uncertainty and sensitivity analyses

### S1 Experimental data

The experimental *in vitro* and *in vivo* data used in our current study are gratefully gathered from a previous study performed by Checkley *et al.* (Checkley et al., 2015). *In vitro* data are listed in Table S1 and *in vivo* data are listed in Tables S2 and S3.

| Time (hours) | Cell count | Std.Dev (count) | $\gamma$H2AX-positive (%) | Std. Dev (%) |
|---|---|---|---|---|
| 0$\mu$M (control) | | | | |
| 2 | 996 | 59.72 | 2.14 | 0.56 |
| 4 | 850 | 62.30 | 2.20 | 0.45 |
| 8 | 1287.5 | 417.59 | 2.90 | 1.22 |
| 16 | 2742.75 | 439.69 | 1.44 | 0.33 |
| 24 | 1857.5 | 409.39 | 1.29 | 0.32 |
| 48 | 3605.25 | 167.38 | 1.93 | 0.44 |
| 72 | 3753 | 311.17 | 1.71 | 0.21 |
| 0.3 $\mu$M | | | | |
| 2 | 1081.88 | 53.63 | 2.93 | 0.59 |
| 4 | 1040.75 | 217.96 | 6.15 | 1.00 |
| 8 | 1447.25 | 392.45 | 7.41 | 1.99 |
| 16 | 2479.5 | 414.02 | 15.68 | 5.56 |
| 24 | 1805.63 | 161.41 | 12.91 | 3.42 |
| 48 | 3497.63 | 385.19 | 11.08 | 4.18 |
| 72 | 3928.25 | 376.08 | 6.57 | 3.30 |
| 1 $\mu$M | | | | |
| 2 | 1129.63 | 58.26 | 17.35 | 3.31 |
| 4 | 1153.63 | 331.31 | 29.12 | 3.47 |
| 8 | 1303.88 | 199.72 | 36.05 | 4.35 |
| 16 | 2420.25 | 744.38 | 38.51 | 9.25 |
| 24 | 1226.38 | 185.58 | 45.01 | 6.01 |
| 48 | 1600.38 | 456.80 | 39.47 | 7.47 |
| 72 | 1612.88 | 540.55 | 33.47 | 5.46 |
| 3 $\mu$M | | | | |
| 2 | 1171.14 | 97.71 | 36.01 | 2.42 |
| 4 | 1291.38 | 567.63 | 46.47 | 4.09 |
| 8 | 1224.63 | 113.30 | 56.72 | 2.62 |
| 16 | 1784.38 | 513.06 | 58.41 | 8.81 |
| 24 | 765.75 | 70.76 | 68.07 | 2.05 |
| 48 | 638.75 | 112.54 | 65.90 | 4.40 |
| 72 | 392.63 | 67.64 | 63.82 | 2.67 |
| 10 $\mu$M | | | | |
| 2 | 1191.13 | 110.15 | 39.38 | 2.62 |
| 4 | 1056.63 | 106.72 | 47.98 | 2.32 |
| 8 | 1113.63 | 144.42 | 59.35 | 1.99 |
| 16 | 1396 | 633.86 | 65.21 | 10.48 |
| 24 | 654.5 | 100.26 | 71.02 | 2.10 |
| 48 | 525.29 | 43.93 | 69.75 | 4.42 |
| 72 | 326.63 | 47.73 | 67.25 | 2.87 |
| 30 $\mu$M | | | | |
| 2 | 1055.13 | 155.16 | 35.37 | 2.21 |
| 4 | 1049.13 | 147.96 | 45.66 | 1.75 |
| 8 | 1228.75 | 211.96 | 51.37 | 1.11 |
| 16 | 1794.88 | 435.42 | 50.35 | 4.19 |
| 24 | 629 | 27.12 | 63.92 | 2.15 |
| 48 | 469.63 | 61.26 | 64.92 | 3.25 |
| 72 | 265.13 | 22.26 | 67.63 | 3.96 |

Table S1: *In vitro* data gathered from a previous study by Checkley *et al.* (Checkley et al., 2015).

| Time (hours) | $\gamma$H2AX-positive (%) | Std. Error (%) |
|---|---|---|
| 0 mg/kg (control) | | |
| 74 | 0.3200 | 0.0320 |
| 80 | 0.2770 | 0.0300 |
| 96 | 0.2970 | 0.0340 |
| 25 mg/kg QD | | |
| 74 | 8.8579 | 0.5364 |
| 80 | 11.3692 | 0.3272 |
| 96 | 12.1945 | 1.0949 |
| 50 mg/kg QD | | |
| 74 | 14.3417 | 0.6278 |
| 80 | 13.8967 | 0.1401 |
| 96 | 17.4986 | 2.7558 |

Table S2: *In vivo* data for DNA damage gathered and adapted from a previous study published by Checkley *et al.* (Checkley et al., 2015).

| Time (hours) | Volume (cm$^3$) | Std. Error (cm$^3$) |
|---|---|---|
| 0 mg/kg (control) | | |
| 168 | 0.3028 | 0.0219 |
| 264 | 0.5189 | 0.0465 |
| 360 | 0.9095 | 0.0934 |
| 456 | 1.3857 | 0.1554 |
| 504 | 1.5646 | 0.1483 |
| 25 mg/kg QD | | |
| 168 | 0.3037 | 0.0342 |
| 264 | 0.4411 | 0.0704 |
| 360 | 0.5617 | 0.0840 |
| 456 | 0.7064 | 0.1221 |
| 504 | 0.8701 | 0.1187 |
| 50 mg/kg QD | | |
| 168 | 0.3106 | 0.0332 |
| 264 | 0.3971 | 0.0768 |
| 360 | 0.4002 | 0.0817 |
| 456 | 0.4783 | 0.0966 |
| 504 | 0.4923 | 0.0846 |

Table S3: *In vivo* data for tumour volume gathered and adapted from a previous study published by Checkley *et al.* (Checkley et al., 2015).

### S2 Model parameterisation

Using a minimal-parameter approach, seven model parameters are calibrated using the *in vitro* data previously produced by Checkley *et al.* (Checkley et al., 2015), as listed in Table 1 in Section 2.7 in the main paper. Parameter sensitivity is explored in the sensitivity analysis (Section 5 of the supplementary material). The calibration process is outlined in Sections S2.1 through to S2.4. The spheroid calibration is described in Section S2.5.

#### S2.1 Cell doubling

In the model, the doubling time of a cell $i$ is denoted $\tau_i$, where $\tau_i$ is stochastically picked from a normal distribution with mean value $\mu$ and standard deviation $\sigma$. Thus $\mu$ corresponds to the average cell doubling time and $\sigma$ corresponds to how synchronised the cells are. If $\sigma$ is zero, then all cells have perfectly synchronised cell cycles and duplicate at the same time. Higher $\sigma$ values achieve less synchronised cell cycles amongst cells and smoother cell count growth curves over time. The control case (i.e. no drug) cell count data is used to estimate $\mu$ and $\sigma$.

By observing the control case cell count data in Table S1, we note that the cell population roughly doubles between 2 and 24 hours (indicating an average cell doubling time, or $\mu$, of approximately 22 hours, which is less than 24 hours). Furthermore, the cell population also roughly doubles between 24 and 48 hours (indicating that $\mu$ is approximately 24 hours). However, in the last 24-hour interval, between 48 and 72 hours, the control population increases by less than 5% (indicating that $\mu$ is more than 24 hours). From these three observations, we choose to make the modelling assumption that the average doubling time for cells should be around 24 hours, and $\sigma$-values in the parameter range [22,26] hours are investigated *in silico*. Due to the synchronised nature of the cell count data, $\sigma$-values between 0 and 2.5 hours were investigated *in silico*, where $\sigma = 0$ h corresponds to completely synchronised cells and $\sigma = 2.5$ h achieves a smooth cell count growth curve. After an iterative process of tuning parameters and running *in silico* experiments, the calibrated values are set to be $\mu = 24$ hours and $\sigma = 0.5$ hours. As discussed in the main article, the ABM can be improved to better fit wet lab data by including variable parameter values or rules, that are updated over time. However, in the current stage of our work, we decided to fix $\mu$ and $\sigma$.

#### S2.2 Cell cycle progression

The *in vitro* data provides information on how many cells are in the damaged S state via the biomarker $\gamma$H2AX. For the control case, the number of $\gamma$H2AX positive cells in our mathematical model depends on two variables: (1) the probability ($\Pi_{D-S}$) that a cell enters the D-S state and (2) the amount of time ($\Theta_{D-S} \cdot \tau_i$) spent in the D-S state prior to repairing. Recall that $\Theta_{D-S}$ is the fraction of a cell's doubling time ($\tau_i$) spent in the D-S state. As a first step, *in silico* experiments are performed in which we find various parameter pairs ($\Pi_{D-S}$, $\Theta_{D-S}$) that agree with the control data. We thereafter note that the *in vitro* drug effect saturates for concentrations 3, 10 and 30 $\mu$M and assume that the maximal dose (30 $\mu$M) yields 100% D-S to S repair inhibition. Thus a second step we test the variable pairs ($\Pi_{D-S}$, $\Theta_{D-S}$) for this 'maximal drug and no repair' scenario *in silico*, and we match these *in silico* results to the 30 $\mu$M *in vitro* data. Here, we only use data from early time points (time $<$ 12 hours) in order to avoid the influence that dying cells have on the data and model outputs. After iterative *in silico* testing, the variable pair ($\Pi_{D-S}$, $\Theta_{D-S}$) that best fits these both extreme cases is $\Pi_{D-S} = 0.75$ and $\Theta_{D-S} = 0.03$. The first extreme case refers to the 'no drug' *in silico* experiment matched to the *in vitro* control data, where we assume that all D-S cells repair to state S. The second extreme case refers to the 'maximum drug' *in silico* experiment matched to the 30$\mu$M control data, where we assume that no D-S cells repair to state S.

#### S2.3 Drug response

Drug effects are modelled using the sigmoid E-max model (Holford, 2017), where the drug effect $E$ is a function of the drug concentration $C$, so that

$$E(C) = E_{max} \cdot \frac{C^\gamma}{EC_{50}^\gamma + C^\gamma},$$

where $E_{max}$ denotes the maximal drug effect. Here we set $E_{max} = 1$ to corresponds to total D-S to S repair inhibition. $EC_{50}$ denotes the drug concentration that achieves half of the maximal drug effect and $\gamma$ is the Hill-coefficient. If drug effect is plotted over time, the $EC_{50}$-value determines the asymptotic behaviour of the effect whilst the $\gamma$-value determines how quickly the asymptotic value is reached.

From the *in vitro* data, we note that the drug concentration 1 $\mu$M achieves roughly half of the total drug effect in terms of $\gamma$H2AX-positive cells. (Note from Table S1 that when the drug concentration is 10 $\mu$M or 30 $\mu$M, the percentage of $\gamma$H2AX-positive cells is roughly 67% at 72 hours, and when the drug concentration is 1 $\mu$M, the percentage of $\gamma$H2AX-positive cells is roughly 33% at 72 hours. Furthermore the lower drug concentration of 0.3 $\mu$M yields a percentage of roughly 7% $\gamma$H2AX-positive at 72 hours, and the higher drug concentration of 3 $\mu$M yields a percentage of roughly 64% $\gamma$H2AX-positive at 72 hours. Consequently, we use 0.3 $\mu$M as a lower bound and 3 $\mu$M as an upper bound for the parameter range in which we seek $EC_{50}$, and $EC_{50}$ values in the range [0.3 $\mu$M, 3 $\mu$M] are investigated with various Hill coefficients to fit *in vitro* data for all (non-control) drug concentrations. In order to avoid the impact that dying cells have on the data used parameterise $EC_{50}$ and $\gamma$, only early *in vitro* data (time $<$ 12 hours) is used to guide the calibration. After iterative *in silico* testing, the best variable pair ($EC_{50}$, $\gamma$) is determined to be $EC_{50} = 1$ $\mu$M and $\gamma = 2$.

#### S2.4 Cell death

In the *in vitro* experiments, cells that are damaged (but not yet dead) are $\gamma$H2AX-positive. In the model, the time it takes between the 'lethal event' (i.e. a cell's failure to repair) and a cell being 'dead' is denoted $T_{L \to D}$ and is matched from the *in vitro* experiment. After noting the asymptotic behaviour of the *in vitro* data, both in terms of cell damage and cell count, we estimate that the rate of cell elimination should roughly correspond to the rate of cell production, and thus $T_{L \to D}$ should be in the same order of magnitude as the doubling time. Consequently, values of $T_{L \to D}$ between 0 and 2 $\tau_i$ are explored *in silico* after which $T_{L \to D} = \tau_i$ is chosen as it best matches the *in vitro* data for all tested (non-control) drug concentrations.

#### S2.5 Spheroid calibration

For the control case, the spheroid model is directly calibrated by the *in vitro* data, and no further calibration is needed. For drug concentrations larger than 0 $\mu$M, we use the *in vivo* data for the highest administered drug dose to calibrate the model in order to disregard details concerning pharmacokinetics and bioavailability. In future work, our model can be integrated with pharmacokinetic modelling techniques.

#### S3 Proliferation of Agents on the Lattice

When an agent divides, it places a daughter agent on the lattice. To achieve an approximately circular population growth on the square lattice, the placement of daughter agents alternates between following the von Neumann (Figure S1(a)) and Moore (Figure S1(a)) convention. The lattice points that are directly adjacent to the lattice point occupied by the parental agent (i.e. the agent that is dividing and thus producing a daughter cell) constitute the 1st order (von Neumann or Moore) neighbourhood of the parental cell, as is illustrated in Figure S1. If a parent is scheduled to place a daughter agent on the lattice according to the von Neumann/Moore convention, then a daughter agent will be placed on a random lattice point in the 1st order von Neumann/Moore neighbourhood. However, if all lattice point in the 1st order neighbourhood (O.N.) are occupied by other agents, the daughter agent is instead placed on a random lattice point in the 2nd O.N. Further, if the 2nd O.N. is also completely occupied by agents, then the daughter agent is placed on a random lattice point in the 3rd O.N., and so on up to the $\nu$th O.N. It is up do the modeller to decide the value of $\nu$, and in this study $\nu$ equals infinity in the *in vitro* monolayer case (with the restriction that agents can not be placed outside the lattice in the *in silico* implementation) and $\tilde{\nu} = 3$ in the spheroid case.

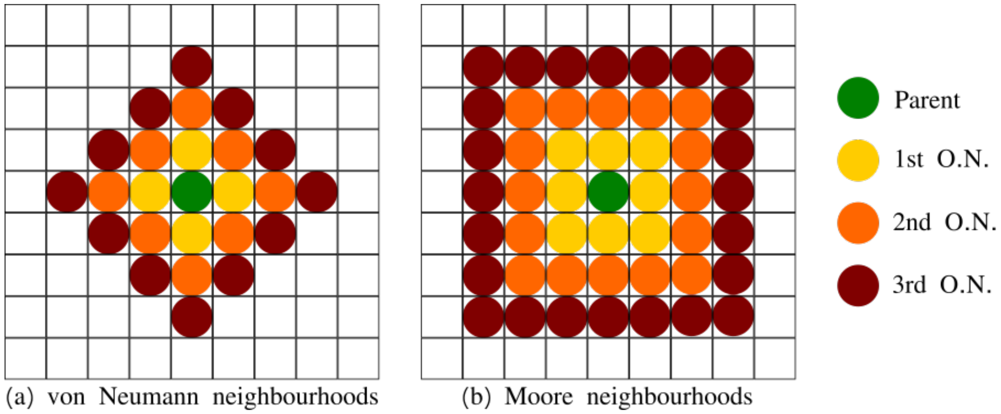


Figure S1: The 1st, 2nd and 3rd order neighbourhood (O.N.) of a parental agent using (a) the von Neumann convention and (b) the Moore convention.

#### S4 Cross-section to tumour spheroid extrapolation

When implementing our mathematical spheroid model, only a central cross-section of the tumour is actually simulated *in silico* and post simulation time this cross-section area (that is approximately circular) is extrapolated to a tumour volume (that is approximately spherical). From the extrapolated tumour spheroid, the two outputs $\tilde{X}_1$ (percentage of $\gamma$H2AX-positive cells) and $\tilde{X}_2$ (tumour volume) are gathered. This is done by using simulated areas to compute the total tumour volume,

$$\tilde{X}_2 = \text{Total Tumour Volume} = \frac{4\pi}{3}\left(\frac{\text{Total Simulated Area}}{\pi}\right)^{3/2}, \tag{1}$$

and the quiescent tumour volume,

$$\text{Quiescent Tumour Volume} = \frac{4\pi}{3}\left(\frac{\text{Quiescent Simulated Area}}{\pi}\right)^{3/2}. \tag{2}$$

From the above, the volume of cycling, or proliferating cells, is obtained by

$$\text{Cycling Tumour Volume} = \text{Total Tumour Volume} - \text{Quiescent Tumour Volume}. \tag{3}$$

Now the output $\tilde{X}_1$ can be computed where,

$$\tilde{X}_1 = \text{Percentage of } \gamma\text{H2AX-positive cells in sphere} = \frac{\text{Number of simulated } \gamma\text{H2AX-positive cells}}{\text{Number of simulated cycling cells}} \times \frac{\text{Cycling Tumour Volume}}{\text{Total Tumour Volume}}. \tag{4}$$

#### S5 A note on simplifying modelling assumptions

Note that an agent in the spheriod setting can, in general, be chosen to comprise either one cancer cell or a group of cancer cells. Note also that modelling the spheroid tumour as a (spatially) two-dimensional disk means that the distribution of nutrients and drugs is modelled across a two-dimensional (rather than a three-dimensional) space. Likewise, the parameter $\tilde{\nu}$, that governs the allowed distance between a parental agent and its daughter agents, only concerns proliferation on a two-dimensional plane. By increasing $\tilde{\nu}$, the tumour will grow quicker and comprise a higher fraction of cycling (to quiescent) agents. Thus both the parameters $\tilde{\nu}$ and $\mu$ influence the rate of tumour growth in the spheroid simulation, and should ideally be fitted to match detailed spheroid data. In the current study, we choose to keep $\mu$ at the *in vitro*-calibrated value and thereafter fit $\tilde{\nu}$ to match the *in vivo* data. A modeller can chose to use a three dimensional spatial domain, and thus explicitly model a tumour spheroid instead of a tumour cross-section. Computational costs, fineness of available data, and the desired level of simulation details should be used to guide the choice of agents and spatial domain.

#### S6 A note on the choice of drug model

We assume all model parameters to be constant in time, including parameters pertaining to the doubling time of the cells. In the monolayer simulation, with some stochastic variability, this complies with Skipper's theoretical law on tumour growth rates which states that, in a tumour growing at a constant exponential rate, the doubling time of the tumour volume is constant over the tumour's lifespan. Further, Skipper's law on tumour regression rates states that the fraction of cells killed by a given drug dose constant is constant (West and Newton, 2017). Although our drug-response model is stochastic, agent-based and concerns a DDR-inhibitor that targets cells in the DNA-damaged state D-S (rather than a general chemotherapy drug that targets cancer cells in other cell cycle states), the probability that a susceptible (D-S) cell is killed by a given drug dose is indeed constant, therefore the percentage of D-S cells killed by a given drug dose is constant (with some stochastic errors/in the large cell population limit). Furthermore, the fraction of cells entering the D-S state is constant and thus our current drug-response model upholds an "essence" of Skipper's tumour regression law. If one instead uses Gompertzian tumour growth models, in which the tumour growth rate varies over the tumour's lifespan, then drug-induced tumour regression can be modelled using the Norton-Simon hypothesis which states that tumor regression is linearly, positively correlated with the tumour growth rate at the time when the treatment is first applied.

#### S7 Residual plots

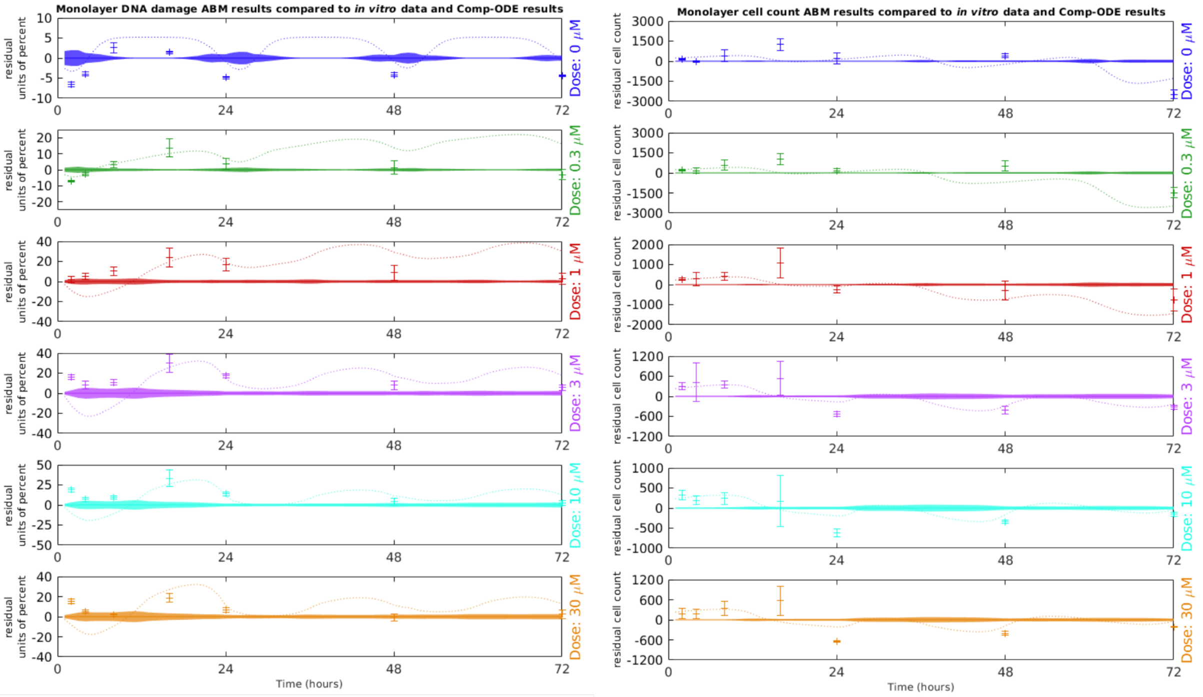


Figure S2: Residual plots for monolayer DNA damage (Left) and cell count (Right) simulation results for different drug doses. Simulation standard deviations are visualised with shaded ribbons. ABM simulation results are compared to *in vitro* data (with standard deviation error bars) (Checkley et al., 2015) and Checkley *et al.*'s compartment-ODE (Comp-ODE) model (Checkley et al., 2015) (dotted lines).

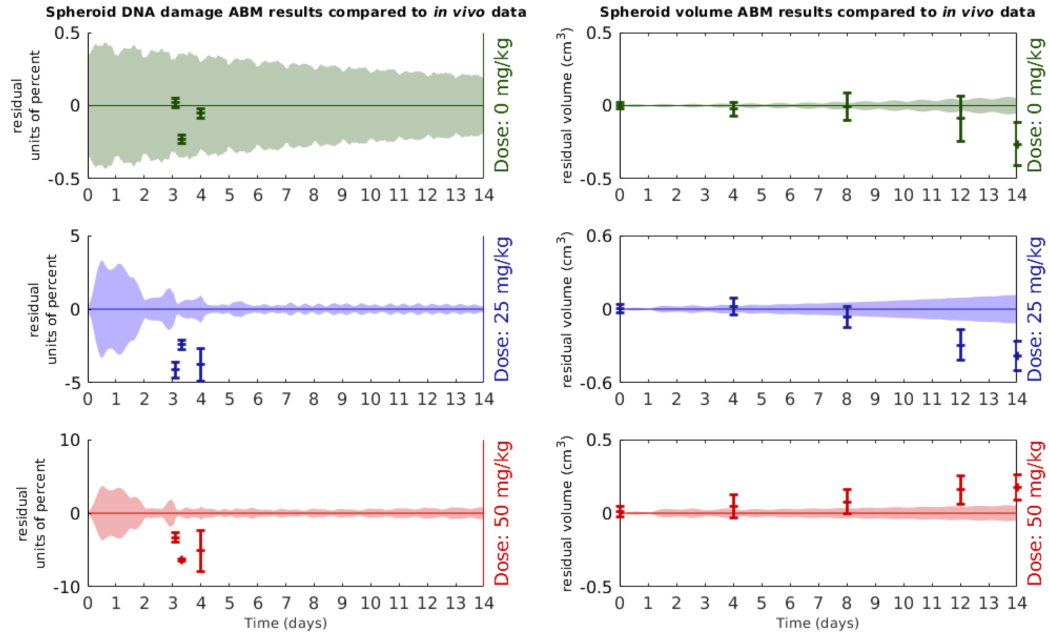


Figure S3: Residual plots where ABM spheroid simulations are compared to *in vivo* xenograft data (Checkley et al., 2015) at different drug doses. Simulation mean values and standard deviations are shown with solid lines and ribbons respectively. Data center points and error bars respectively show *in vivo* mean values and standard errors.

### S8 Spheroid waterfall plot

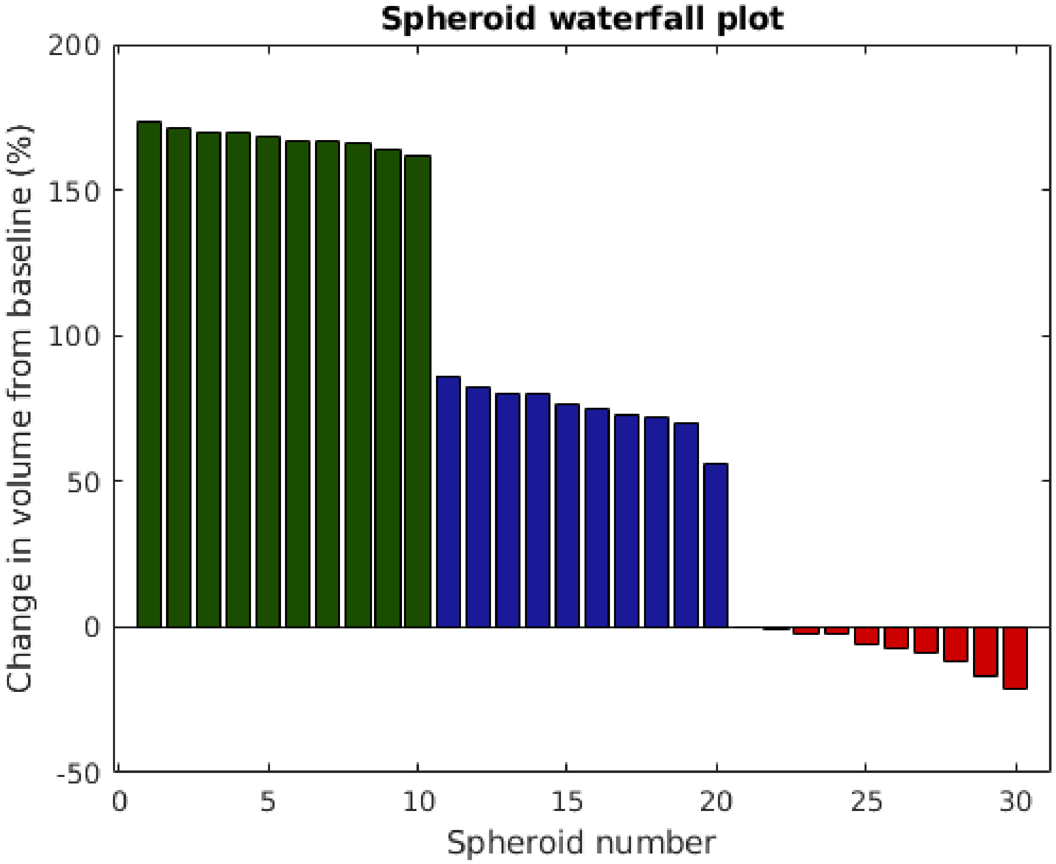


Figure S4: Waterfall plots showing the change in tumour volume at day 7 from the baseline (the tumour volume when treatment commences). Spheroids are subjected to drugs once daily at doses 25 mg/kg (blue bars) or 50 mg/kg (red bars). Green bars correspond to control cases (no drug).

#### S9 Uncertainty and Sensitivity Analyses

To evaluate the monolayer *in silico* findings obtained in our *in vitro* study, three uncertainty and sensitivity analyses techniques are performed. The three techniques are namely: (1) Consistency Analysis, which is used to determine how many *in silico* runs should be performed before defining results in terms of statistical metrics in order to mitigate uncertainty originating from intrinsic model stochasticity, (2) Robustness Analysis, which investigates model sensitivity to local parameter perturbations and (3) Latin Hypercube Analysis, which investigates model sensitivity to global parameter perturbations. These techniques are thoroughly described in a review/methodology paper (Hamis et al., 2020) which provides background information concerning the origin of these techniques as well as detailed information on how to implement them. To perform uncertainty and sensitivity analyses we need to specify a set of inputs and outputs. Here, the output variables are $X_1$: the percentage of $\gamma$H2AX-positive (i.e. damaged) cells at the end time of the experiment (72 hours), and $X_2$: the cell count (i.e. the number of non-dead cells) at the end of the experiment. The input variables are the seven model parameters listed in Table 1, in the main article, that we calibrate using *in vitro* data. These inputs are namely $\mu$, $\sigma$, $\Pi_{D-S}$, $\Theta_{D-S}$, $EC_{50}$, $\gamma$ and $T_{L\to D}$.

##### S9.1 Consistency Analysis

Results from the Consistency Analysis are provided in Figures S5, S6, S7, S8, S9 which show the $\hat{A}$-measures, in both computed and scaled forms, for the distribution sizes $n = 1, 5, 50, 100, 300$ respectively. By observing Figures S5 through to S9, it is clear that the statistical significance decreases with increasing distribution size $n$, as is shown in Figure S10 and Table S9.1 which show the maximal scaled $\hat{A}$-values for all tested distribution sizes. These results demonstrate that the distribution size $n = 100$ is the smallest tested distribution size that yields a small statistical significance (i.e. a maximum scaled $\hat{A}$-value smaller than 0.56) for both regarded output variables $X_1$ and $X_2$. From this we decide to base every *in silico* result (here in terms of mean values and standard deviations) on 100 simulation runs.

| output \ distribution size | n=1 | n=5 | n=50 | n=100 | n=300 |
|---|---|---|---|---|---|
| $X_1$ | 1 | 0.92 | 0.61 | 0.55 | 0.54 |
| $X_2$ | 1 | 0.84 | 0.59 | 0.55 | 0.54 |

Table S4: Maximal scaled $\hat{A}$-values produced in the Consistency Analysis for various distribution sixes $n$. The output variables are $X_1$, corresponding to the percentage of $\gamma$H2AX positive (i.e. damaged) cells, and $X_2$, corresponding to the cell count.

#### S9.2 Robustness Analysis

We use Robustness Analysis to investigate how sensitive the output is to *local* parameter perturbations, that is to say when input parameters are varied one at a time. Figures S11, S12, S13, S14, S15, S16, S17 provide boxplots and $\hat{A}$-measures that demonstrate the effect that local perturbations of the input variables $\mu$, $\sigma$, $\Pi_{D-s}$, $\Theta_{D-S}$, $EC_{50}$, $\gamma$ and $T_{L\to D}$ respectively have on the output variables $X_1$ and $X_2$. Key findings are listed below, discussing the impact of one input parameter at a time.

- Remarks regarding input parameter $\mu$: Figure S11 shows that, for small parameter perturbations, increasing the average doubling times of cells, $\mu$, overall decreases the percentage of $\gamma$H2AX positive cells and increases the cell count, however this decrease/increase is not linear. This indicates that the results of the *in vitro* monolayer simulation (and of the *in vitro* experiment nonetheless) are sensitive to the *timing* of the drug administration. In other words, Robustness Analyses demonstrates that treatment responses depend on how many cells are in the susceptible cell-cycle state at time of drug administration.

- Remarks regarding input parameter $\sigma$: Figure S12 demonstrates that the level of cell cycle synchronisation amongst cells, quantified by the input $\sigma$, affects *in silico* outputs for small parameter perturbations. The results indicate that for highly asynchronised cells (i.e. high $\sigma$-values) the smoother growth curves yield higher cell counts at certain time-points (such as the end time 72 hours) and a lower percentage of $\gamma$H2AX-positive cells. As discussed in the remark above, the timing between cell cycles and drug administration affect treatment responses.

- Remarks regarding input parameter $\Pi_{D-S}$: Figure S13 illustrates that increasing the probability that a cell enters the damaged S state, i.e. the variable $\Pi_{D-S}$, increases the percentage of $\gamma$H2AX cells and decreases the cell count, as expected.

- Remarks regarding input parameter $\Theta_{D-S}$: Figure S14 shows how the amount of time that damaged cells spend in the D-S state before attempting to repair, and thus the $\Theta_{D-S}$-value, affects the output. Results show that the percentage of $\gamma$H2AX positive cells increases with increasing values of $\Theta_{D-S}$, as more damaged cells will accumulate in the D-S state. However, this does not affect the probability of cells repairing, so the cell count is not as sensitive to small perturbations of $\Theta_{D-S}$. The value of $\Theta D - S$ implicitly affects the measured cell count at the end time of the experiment as a decreased/increased $\Theta D - S$-value yields a slightly decreased/increased time lag between a cell entering the D-S state and dying.

- Remarks regarding input parameter $EC_{50}$: Figure S15 demonstrates that output variables are highly sensitive to perturbations of $EC_{50}$. Increasing $EC_{50}$ results in a higher percentage of $\gamma$H2AX positive cells and a lower cell count. Thus the input parameter $EC_{50}$ should be regarded as a highly influential on quantitative results.

- Remarks regarding input parameter $\gamma$: Figure S16 illustrates that output variables measured at the end time of the experiment are not very sensitive to small perturbations of $\gamma$.

This can be understood as the $\gamma$ parameter inherently corresponds to 'how quickly' a drug achieves asymptotic behaviour in the $E_{max}$ model, the model used in our mathematical framework to express cellular drug response.

- Remarks regarding input parameter $T_{L\rightarrow}$: Figure S17 shows how output variables change as a result of perturbations to the input variable $T_{L\rightarrow}$, that describes how long it takes for a cell that has failed to repair to die (i.e. how long a cell with a 'death-sentence' is picked up as $\gamma$H2AX positive in the *in vitro* experiment). Results show that both the percentage of $\gamma$H2AX positive cells and the cell count increases with increasing values of $T_{L\rightarrow}$, as dying cells will categorised as $\gamma$H2AX positive longer before being categorised as dead. When calibrating the model, we avoid the effect of this input parameter by only regarding *in vitro* data at time points that are early enough to correspond to systems with no (or a negligible amount of) dead cells.

#### S9.3 Latin Hypercube Analysis

Latin Hypercube Analysis is here used to investigate how sensitive output responses are to *global* parameter perturbations. We here investigate parameter values within ranges that we consider to be 'plausible' from the calibration process and the Robustness Analysis. Figures S18, S19, S20, S21, S22, S23, S24 provide scatter-plots that demonstrate correlations between the output variables $X_1$ and $X_2$ and the input variables $\mu$, $\sigma$, $\Pi_{D-s}$, $\Theta_{D-S}$, $EC_{50}$, $\gamma$ and $T_{L\rightarrow D}$ respectively. The Pearson Product Moment Correlation Coefficients between the various input-output pairs are listed in Table S9.3. To determine threshold values for correlation coefficient descriptors, we compromise between suggested values by other authors (Hamis et al., 2020), and take into account the fact that we are only regarding parameter values within 'plausible' ranges. With this as a guide, we here decide that our obtained correlation coefficients with a magnitude in [0,0.12] corresponds to the linear input-output relationship being 'negligible', [0.19, 0.35] 'weak', [0.48,0.59] 'moderate' and 0.84 'strong'. Key findings from the Latin Hypercube Analysis are listed below, where the impact of one input parameter is discussed one at a time.

- Remarks regarding input parameter $\mu$: Figure S18 and the first column in Table S9.3 show that, for the allowed parameter range, $\mu$ and $X_1$ are moderately, negatively correlated as the correlation coefficient is -0.48 and the scatterplot displays an overall trend of the output ($X_1$) decreasing with increasing values of the input $\mu$. The relationship between $\mu$ and the other output variable $X_2$ is, on the other hand, negligible. We explain this by the fact that treatment responses are sensitive to the timing of the drug administration, but there is a time-lag $T_{L\rightarrow D}$ between a cell's lethal event (failure to repair) and its death. As damaged (but not dead) cells are included in the cell count, the ($\mu$,$X_2$)-relationship is more weakly linearly correlated than the ($\mu$,$X_1$)-relationship.

- Remarks regarding input parameter $\sigma$: Figure S19 and the second column in Table S9.3 demonstrate that the linear relationships between input variable $\sigma$ and the output variables $X_1$ and $X_2$ are both negligible, within the regarded input parameter value range.

- Remarks regarding input parameter $\Pi_{D-S}$: Figure S20 and the third column in Table S9.3 indicate that the relationships between the input variable $\Pi_{D-S}$ and the output variables

$X_1$ and $X_2$ are, respectively, positively and negatively weakly linearly correlated. This agrees with the intuitive notion that if the probability that a cell enters the D-S state increases, cell damage ($X_1$) increases whilst the cell count ($X_2$) decreases.

- Remarks regarding input parameter $\Theta_{D-S}$: Figure S21 and the fourth column in Table S9.3 show that the input variable $\Theta_{D-S}$ is has a negligible linear correlation with the output variables $X_1$ and $X_2$.

- Remarks regarding input parameter $EC_{50}$: Figure S22 and the fifth column in Table S9.3 demonstrate that the input variable $EC_{50}$ impacts the output responses more than do other input variables, within the regarded ranges for input variables. $EC_{50}$ is negatively, moderately linearly correlated with $X_1$ and $EC_{50}$ is strongly, positively linearly correlated with $X_2$. These relationships are visually apparent in the regarded scatterplots.

- Remarks regarding input parameter $\gamma$: Figure S23 and the sixth column in Table S9.3 indicate negligible linear correlations between the input parameter $\gamma$ and both output variables $X_1$ and $X_2$.

- Remarks regarding input parameter $T_{L\to D}$: Figure S24 and the last column in Table S9.3 demonstrate that the input variable $T_{L\to D}$ is positively, weakly, linearly correlated with the output $X_1$, whilst the linear correlation between $T_{L\to D}$ and $X_2$ is negligible.

| input / output | $\mu$ | $\sigma$ | $\Pi_{D-S}$ | $\Theta_{D-S}$ | $EC_{50}$ | $\gamma$ | $T_{L\to D}$ |
|---|---|---|---|---|---|---|---|
| $X_1$ | -0.48 | 0.06 | 0.19 | 0.06 | -0.59 | 0.05 | 0.35 |
| $X_2$ | 0.12 | 0.01 | -0.24 | -0.02 | 0.84 | 0.12 | 0.00 |

Table S5: Pearson Product Moment Correlation Coefficients between input and output variables obtained in the Latin Hypercube Analysis.

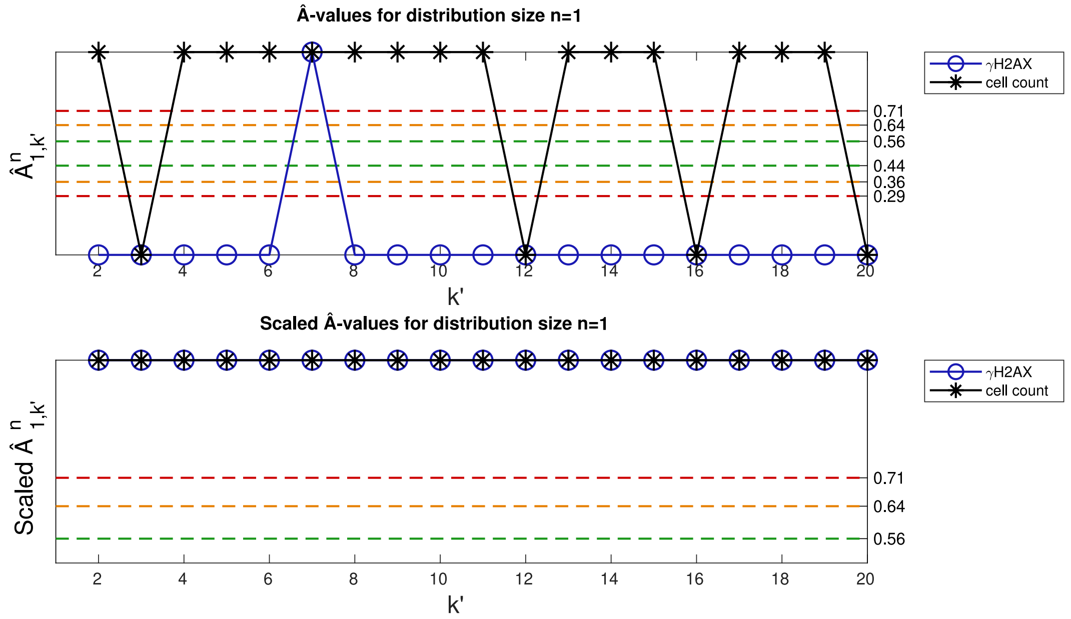


Figure S5: Consistency Analysis. $\hat{A}$-values in initial (top) and scaled (bottom) form for distribution size $n = 1$.

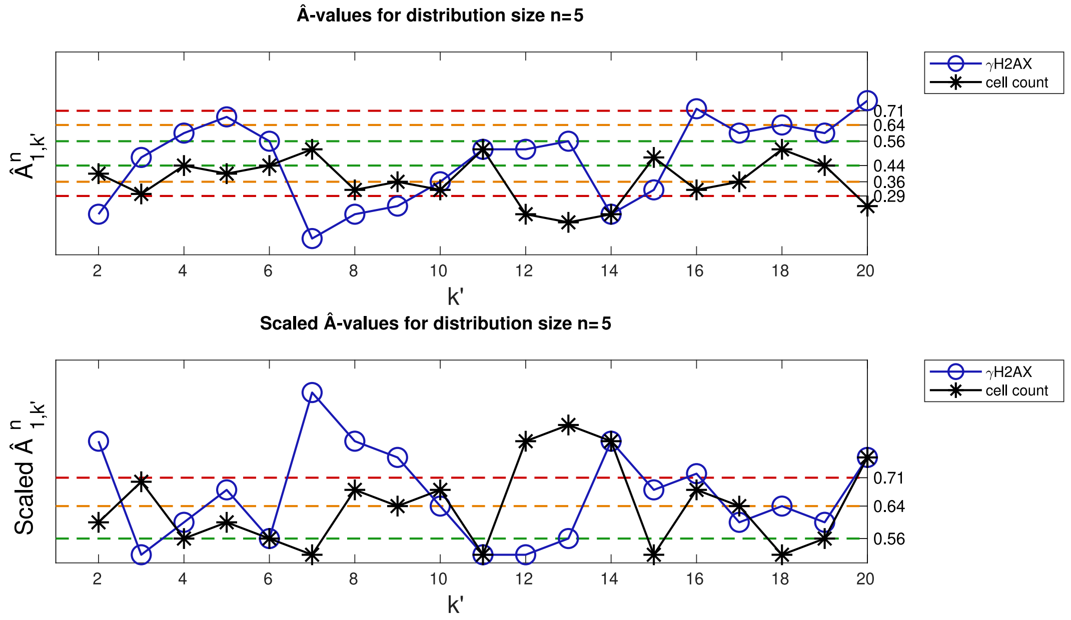


Figure S6: Consistency Analysis. $\hat{A}$-values in initial (top) and scaled (bottom) form for distribution size $n = 5$.

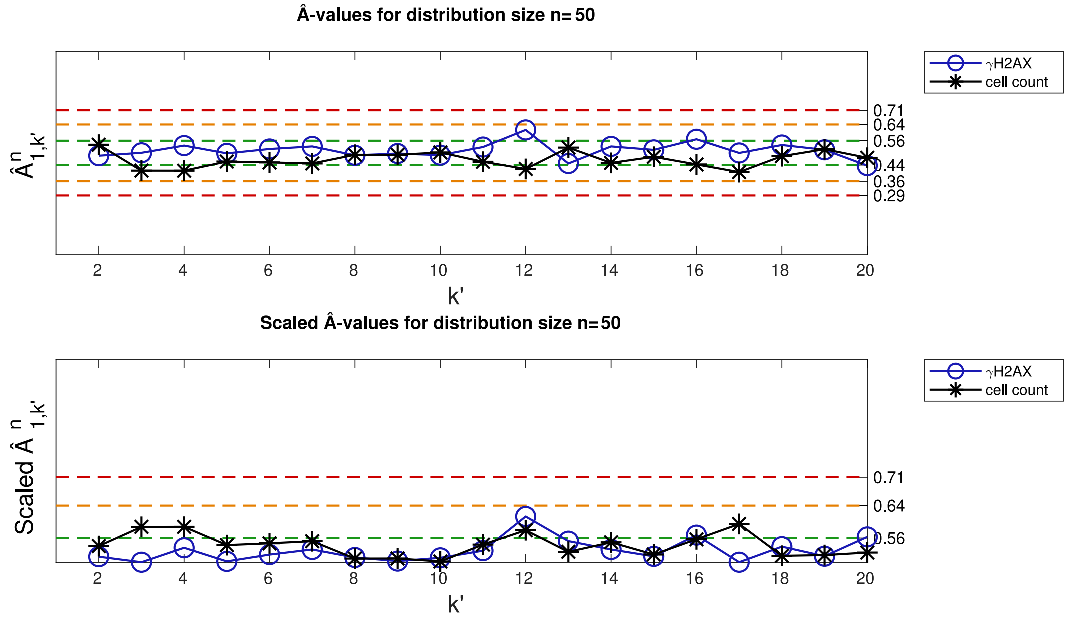


Figure S7: Consistency Analysis. $\hat{A}$-values in initial (top) and scaled (bottom) form for distribution size $n = 50$.

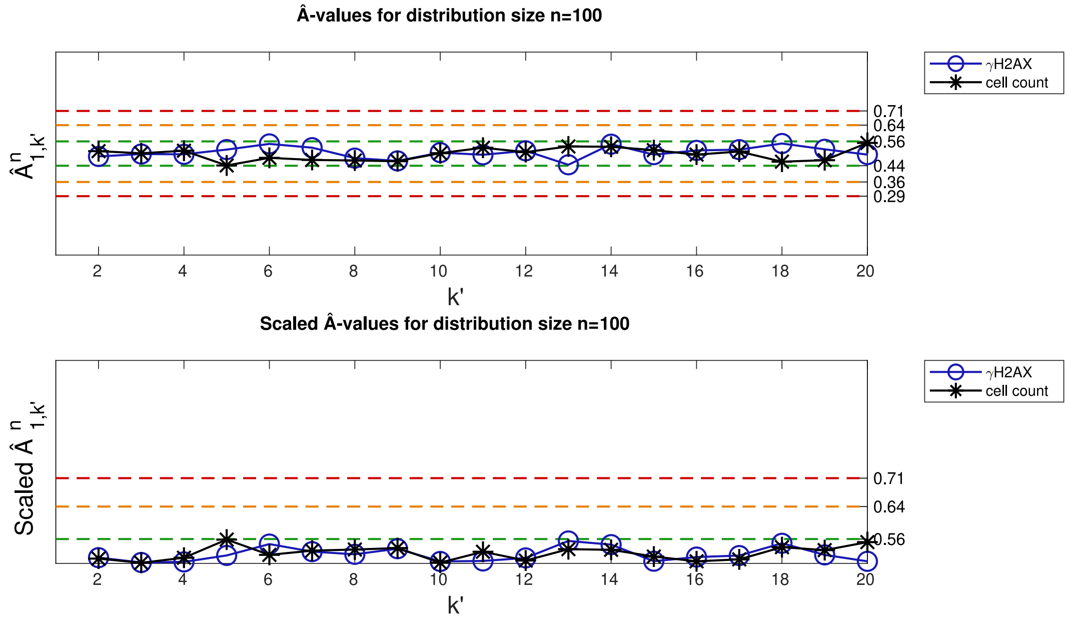


Figure S8: Consistency Analysis. $\hat{A}$-values in initial (top) and scaled (bottom) form for distribution size $n = 100$.

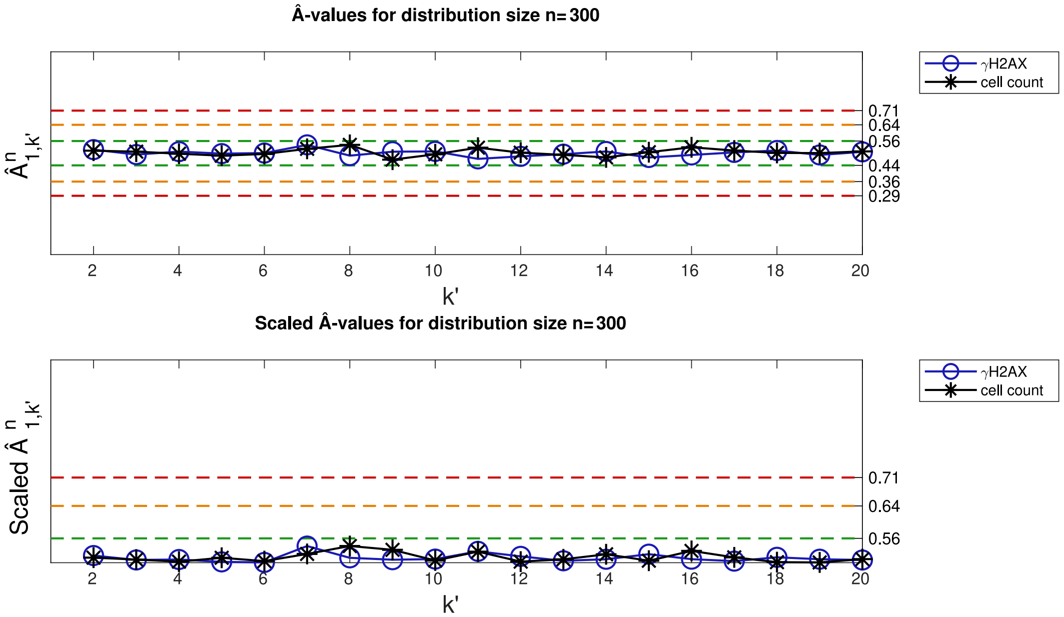


Figure S9: Consistency Analysis. $\hat{A}$-values in initial (top) and scaled (bottom) form for distribution size $n = 300$.

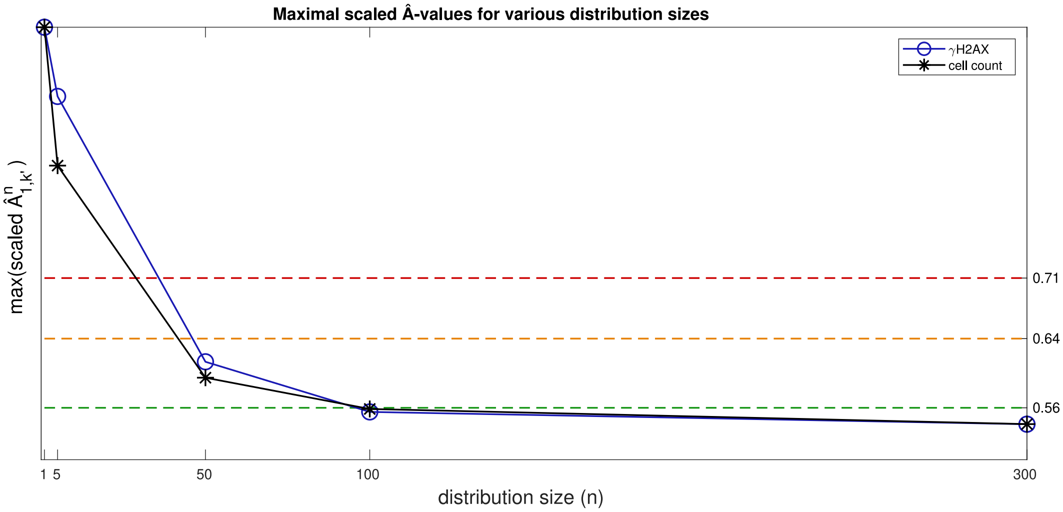


Figure S10: Consistency Analysis. Scaled $\hat{A}$-values for various distribution sizes tested in the Consistency Analysis.

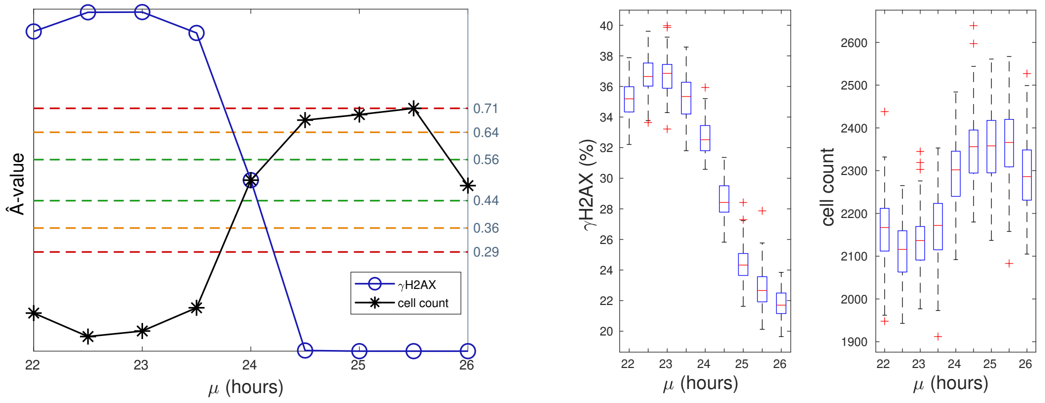


Figure S11: Robustness Analysis. Left: Output responses, in terms of percentage of $\gamma$H2AX positive (i.e. damaged) cells, and cell count as a result of perturbations to the input variable $\mu$. Right: Maximal $\hat{A}$-values resulting from comparisons between distributions with perturbed data and a distribution with calibrated (unperturbed) data.

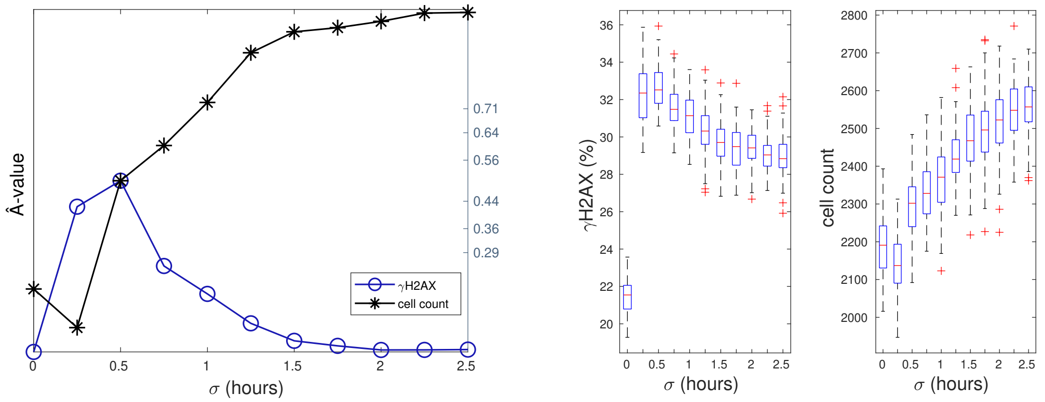


Figure S12: Robustness Analysis. Left: Output responses, in terms of percentage of $\gamma$H2AX positive (i.e. damaged) cells, and cell count as a result of perturbations to the input variable $\sigma$. Right: Maximal $\hat{A}$-values resulting from comparisons between distributions with perturbed data and a distribution with calibrated (unperturbed) data.

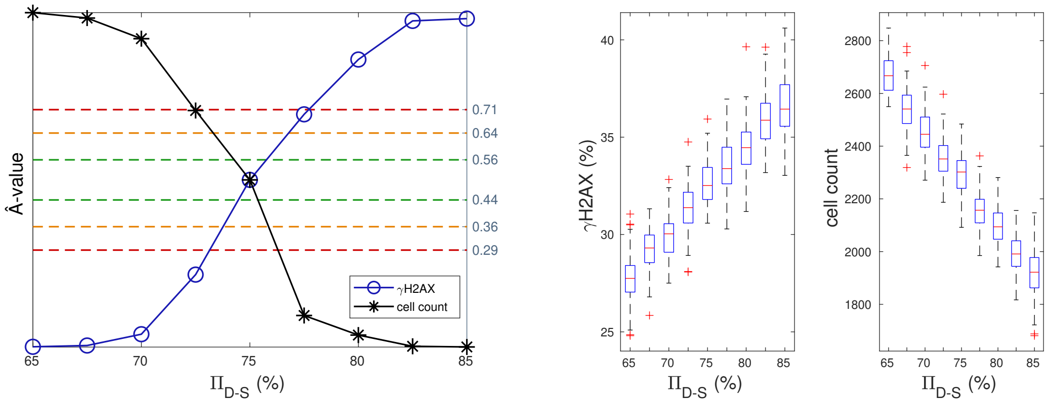


Figure S13: Robustness Analysis. Left: Output responses, in terms of percentage of $\gamma$H2AX positive (i.e. damaged) cells, and cell count as a result of perturbations to the input variable $\Pi_{D-s}$. Right: Maximal $\hat{A}$-values resulting from comparisons between distributions with perturbed data and a distribution with calibrated (unperturbed) data.

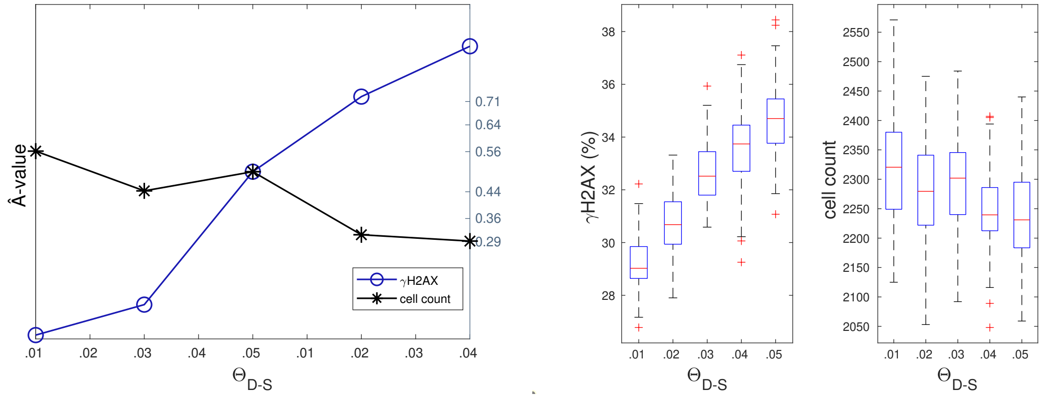


Figure S14: Robustness Analysis. Left: Output responses, in terms of percentage of $\gamma$H2AX positive (i.e. damaged) cells, and cell count as a result of perturbations to the input variable $\Theta_{D-S}$. Right: Maximal $\hat{A}$-values resulting from comparisons between distributions with perturbed data and a distribution with calibrated (unperturbed) data.

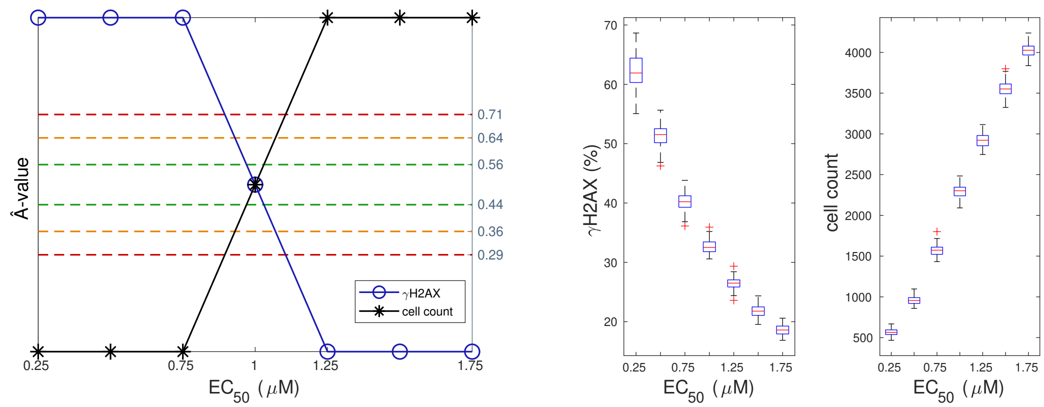


Figure S15: Robustness Analysis. Left: Output responses, in terms of percentage of $\gamma$H2AX positive (i.e. damaged) cells, and cell count as a result of perturbations to the input variable $EC_{50}$. Right: Maximal $\hat{A}$-values resulting from comparisons between distributions with perturbed data and a distribution with calibrated (unperturbed) data.

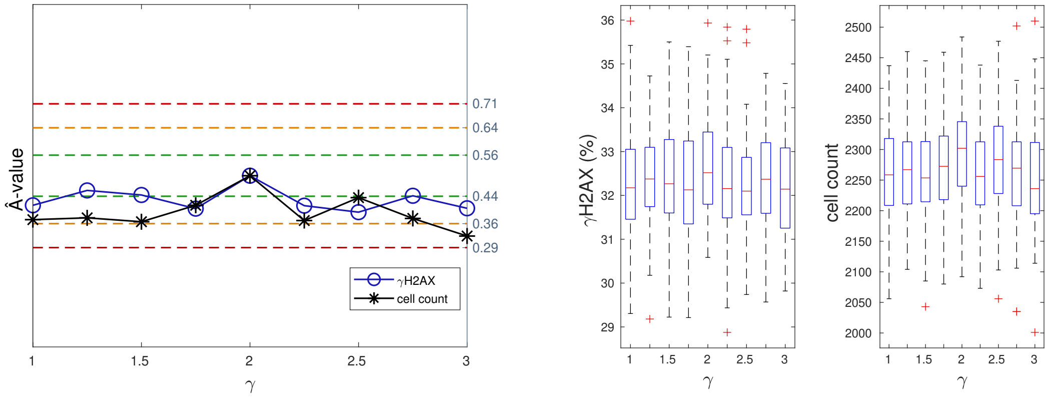


Figure S16: Robustness Analysis. Left: Output responses, in terms of percentage of $\gamma$H2AX positive (i.e. damaged) cells, and cell count as a result of perturbations to the input variable $\gamma$. Right: Maximal $\hat{A}$-values resulting from comparisons between distributions with perturbed data and a distribution with calibrated (unperturbed) data.

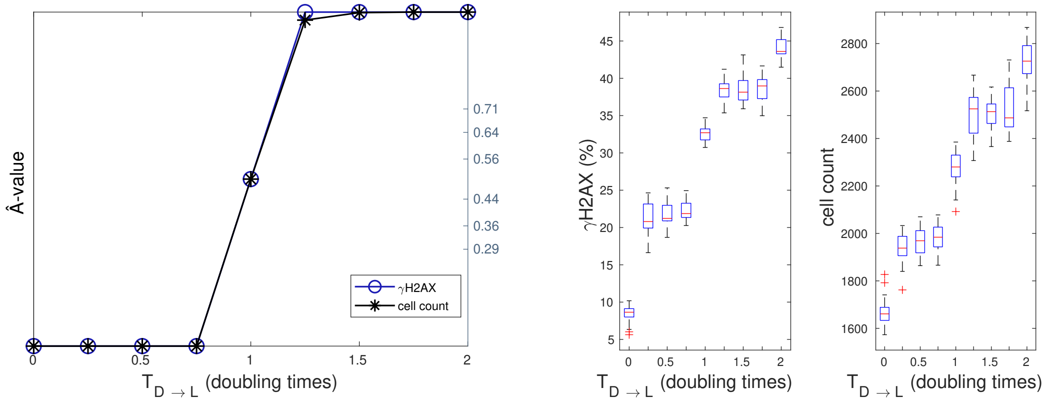


Figure S17: Robustness Analysis. Left: Output responses, in terms of percentage of $\gamma$H2AX positive (i.e. damaged) cells, and cell count as a result of perturbations to the input variable $T_{D \to L}$. Right: Maximal $\hat{A}$-values resulting from comparisons between distributions with perturbed data and a distribution with calibrated (unperturbed) data.

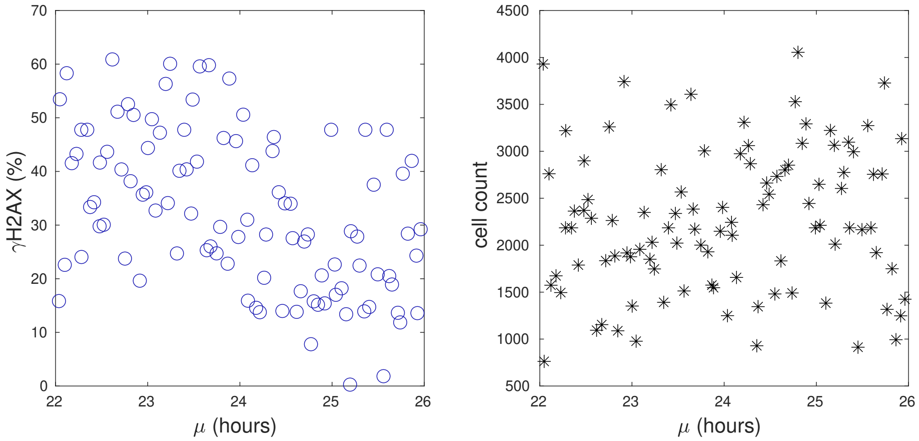


Figure S18: Latin Hypercube Analysis. Outputs in terms of $\gamma$H2AX positive cells (left) and number of viable cells (right) when global parameter perturbations are performed. The scatterplots show the correlation between outputs and the input variable $\mu$.

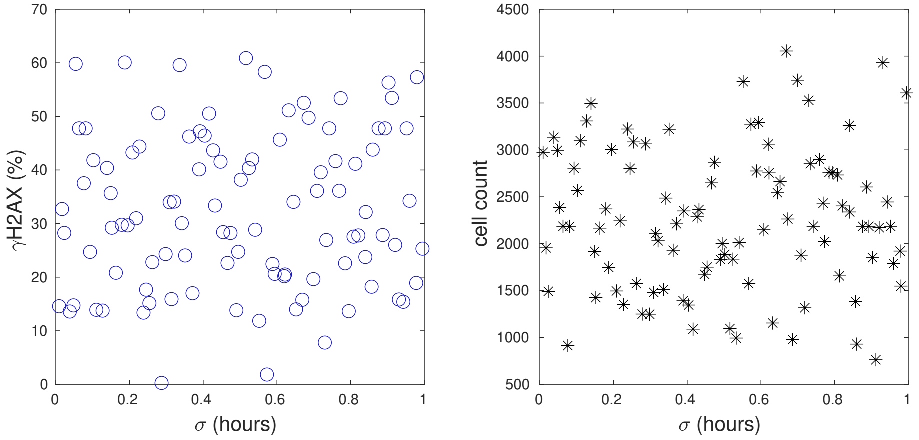


Figure S19: Latin Hypercube Analysis. Outputs in terms of $\gamma$H2AX positive cells (left) and number of viable cells (right) when global parameter perturbations are performed. The scatter-plots show the correlation between outputs and the input variable $\sigma$.

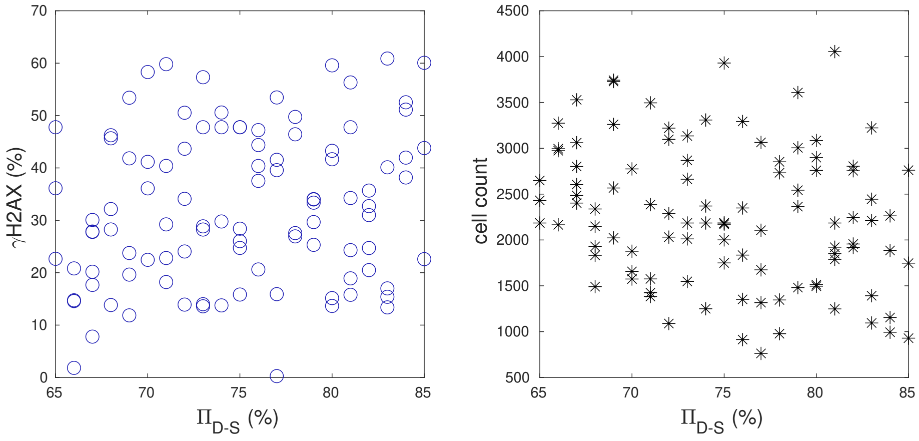


Figure S20: Latin Hypercube Analysis. Outputs in terms of $\gamma$H2AX positive cells (left) and number of viable cells (right) when global parameter perturbations are performed. The scatter-plots show the correlation between outputs and the input variable $\Pi_{D-S}$.

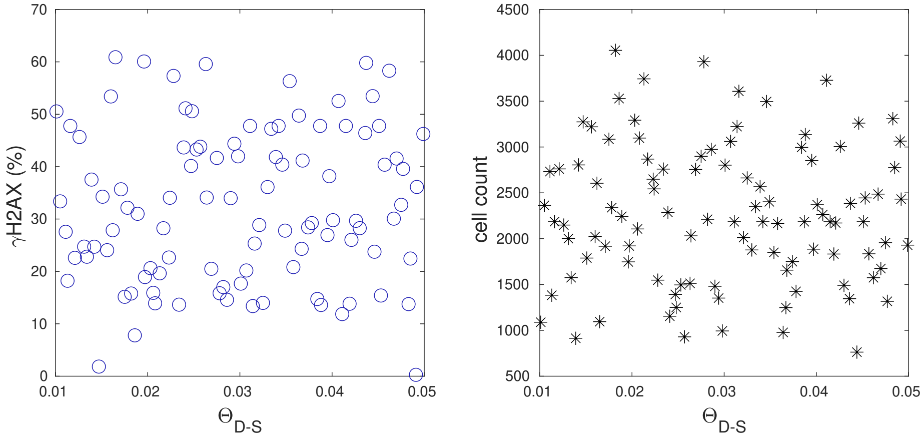


Figure S21: Latin Hypercube Analysis. Outputs in terms of $\gamma$H2AX positive cells (left) and number of viable cells (right) when global parameter perturbations are performed. The scatterplots show the correlation between outputs and the input variable $\Theta_{D-S}$.

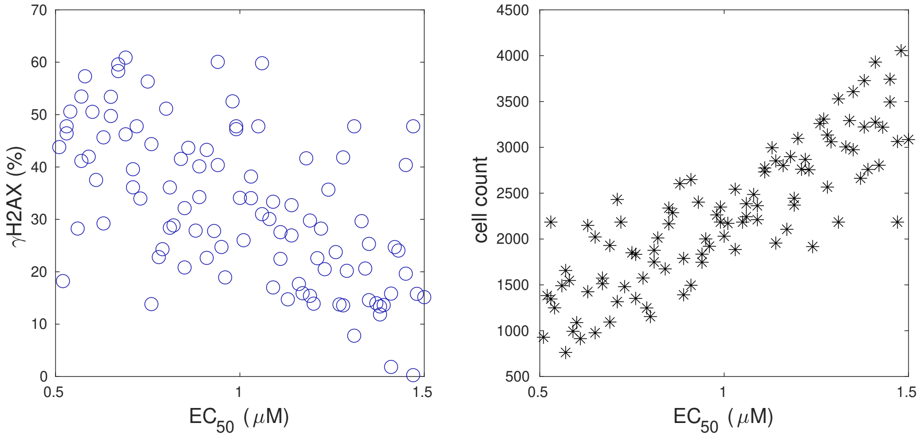


Figure S22: Latin Hypercube Analysis. Outputs in terms of $\gamma$H2AX positive cells (left) and number of viable cells (right) when global parameter perturbations are performed. The scatterplots show the correlation between outputs and the input variable $EC_{50}$.

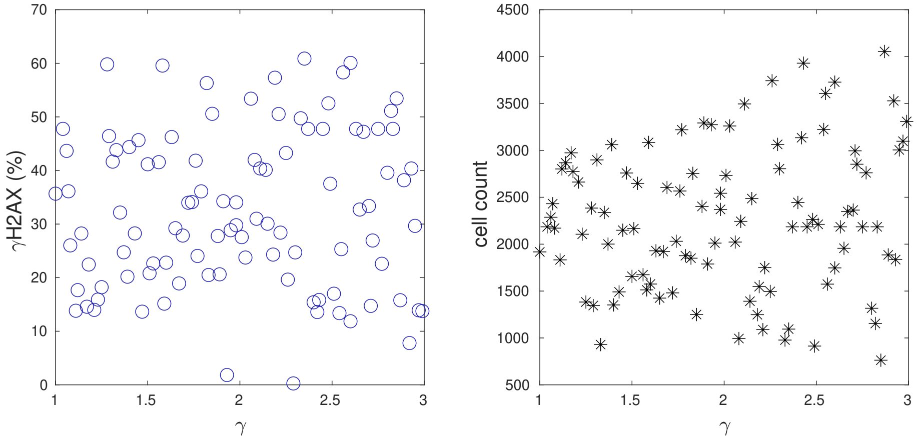


Figure S23: Latin Hypercube Analysis. Outputs in terms of $\gamma$H2AX positive cells (left) and number of viable cells (right) when global parameter perturbations are performed. The scatterplots show the correlation between outputs and the input variable $\gamma$.

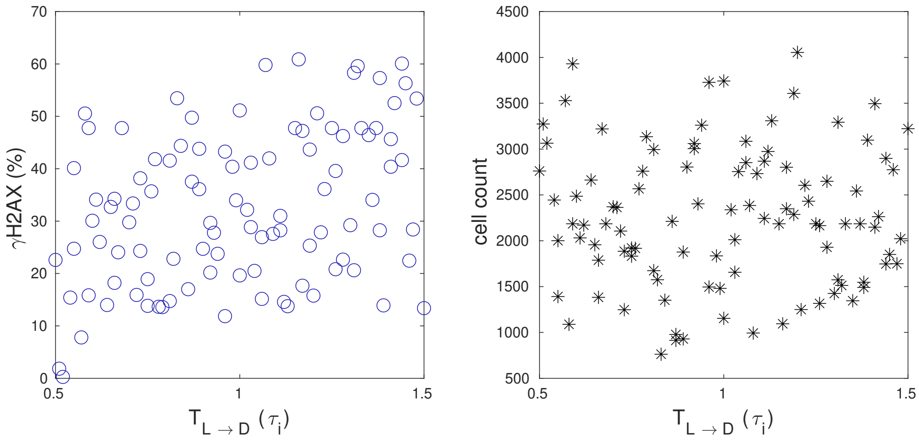


Figure S24: Latin Hypercube Analysis. Outputs in terms of $\gamma$H2AX positive cells (left) and number of viable cells (right) when global parameter perturbations are performed. The scatterplots show the correlation between outputs and the input variable $T_{L\to D}$.
